# Intrinsic errors in mitochondrial translation trigger a decline in cell fitness

**DOI:** 10.1101/2025.10.13.682189

**Authors:** Güleycan Lutfullahoglu Bal, Kah Ying Ng, Eli Berzell, Ani Akpinar, Alex E. Ekvik, Lidiia Koludarova, Seyedehshima Naddafi, Clemens Raths, Tania Vane-Tempest, Jordyn J. VanPortfliet, Sarah O’Keefe, Arafath K. Najumudeen, Tuula A. Nyman, James B. Stewart, A. Phillip West, Brendan J. Battersby

**Author notes:** Department of Nutritional Sciences and Toxicology, University of California, Berkeley, Berkeley, CA 94720, USA.

## Abstract

Defects in the faithful expression of the human mitochondrial genome underlies disease states, from rare inherited disorders to common pathologies and the aging process itself. The ensuing decrease in the capacity for oxidative phosphorylation alone cannot account for the phenotype complexity associated with disease. Here, we address how aberrations in mitochondrial nascent chain synthesis *per se* exert a decline in cell fitness using a classic model of mitochondrial induced premature aging. We identify how intrinsic errors during mitochondrial nascent chain synthesis destabilize organelle gene expression, triggering intracellular stress responses that rewire cellular metabolism and cytokine secretion. Further, we show how these mechanisms extend to pathogenic variants associated with inherited human disorders. Together, our findings reveal how aberrations in mitochondrial protein synthesis can sensitize a cell to metabolic challenges associated with disease and pathogen infection independent of oxidative phosphorylation.

**Teaser/One-Sentence Summary:** Aberrations in mitochondrial translation elongation trigger activation of intracellular stress responses associated with disease and aging.

---

Aging is defined by mitochondrial dysfunction. The progressive accumulation of mitochondrial DNA (mtDNA) mutations and their expression is associated with an age-related decline in cell fitness (*1–3*). In metazoans, mtDNA is a small multi-copy genome encoding only 13 essential proteins of the oxidative phosphorylation (OXPHOS) complexes (*4*). The biogenesis of each of these proteins requires co-translational insertion into the mitochondrial inner membrane (*5*). As seen in aging, translation of mitochondrial mRNAs encoding an elevated number of mutations leads to an increased rate of nascent chain turnover (*6*). This impairs assembly of the OXPHOS complexes. A classical view is that quantitative defects in OXPHOS capacity is the catalyst for the aging phenotypes (*6*, *7*). However, inherited mitochondrial disorders manifest with exceptionally diverse clinical presentations that differ in their severity, age of onset and tissue-specificity (*8*). Thus, the impact of mitochondrial dysfunction on cell fitness is far more complex and other, OXPHOS independent, factors must confer important molecular contributors to pathology.

Mitochondria lack a physical membrane barrier to separate transcription, RNA processing and translation. Hence, aberrations in gene expression exert a direct effect on mitochondrial ribosomes and the organelle inner membrane. Recent studies reveal that mitochondrial mRNAs with aberrant 3’ ends are constantly generated and that there is no safeguard mechanism to prevent translation initiation on these transcripts, even in healthy cells and tissues (*9*, *10*). One notable class of such transcripts are fusion open-reading frames (ORFs) (*9*, *10*). Translation of these fusion ORFs will induce nascent chain misfolding and require rapid proteolytic degradation by the membrane anchored AAA metalloprotease complexes composed of the AFG3L2 subunit (*11–13*). This nascent chain quality control complex is paramount for maintaining mitochondrial gene expression, membrane morphology, metabolic function and cell fitness (*11–19*).

We hypothesized that the inherent error rates in mitochondrial gene expression during nascent chain synthesis, folding and membrane insertion would be compounded by the increased expression of mtDNA mutations associated with aging. Further, we posited that it is the translation of mRNAs encoding these accumulated deleterious mutations on mitochondrial ribosomes *per se* that can act as a negative driver on cell fitness. Thus, by inhibiting mitochondrial protein synthesis, we sought to identify aging associated cellular phenotypes that are independent of OXPHOS dysfunction.

To experimentally test our hypothesis requires the integration of several key concepts. First, we harnessed mouse embryonic fibroblasts (MEFs) derived from the *Polg ^D257A^*mtDNA mutator mouse, a classic model of mammalian aging (*1*, *2*). These mice encode a single point mutation in the proof-reading domain of the nuclear-encoded mitochondrial DNA polymerase (*Polg*), generating random mutations across the mitochondrial genome with each round of mtDNA replication (*1*, *2*, *20*, *21*). We subjected these MEFs to long-term culture (Fig. 1A). Next, we exploited the well-characterized antibiotic chloramphenicol to inhibit mitochondrial protein synthesis. This specific inhibitor of mitochondrial translation elongation acts only when the amino acid in the −1 position of the nascent polypeptide chain is an alanine, serine or threonine (*22*). These three residues are highly enriched within the 13 mtDNA-encoded proteins (fig. S1A). Consequently, long-term culture of *Polg ^D257A^* MEFs with chloramphenicol prevents the synthesis of these 13 nascent chains, blocking OXPHOS complex assembly and function. Despite this severe bioenergetic challenge to the cell, mitotic mammalian cell types can be extensively cultured with chloramphenicol when supplemented with uridine (fig. S1C) (*11*, *12*, *14*). Inclusion of this nucleoside in the culture media bypasses the need of the respiratory chain in *de novo* pyrimidine biosynthesis (*23*). Phenotypically, chloramphenicol leads to remodeling of the cristae inner membrane ultrastructure from the lack of OXPHOS complexes (*12*), but exerts no effect on the overall reticular membrane morphology of the organelle (fig. S1D). Together, our experimental design allows us to investigate the specific contribution of mitochondrial protein synthesis itself to the age-related decline in cell fitness associated with the expression of mtDNA mutations.

**Fig. 1.**
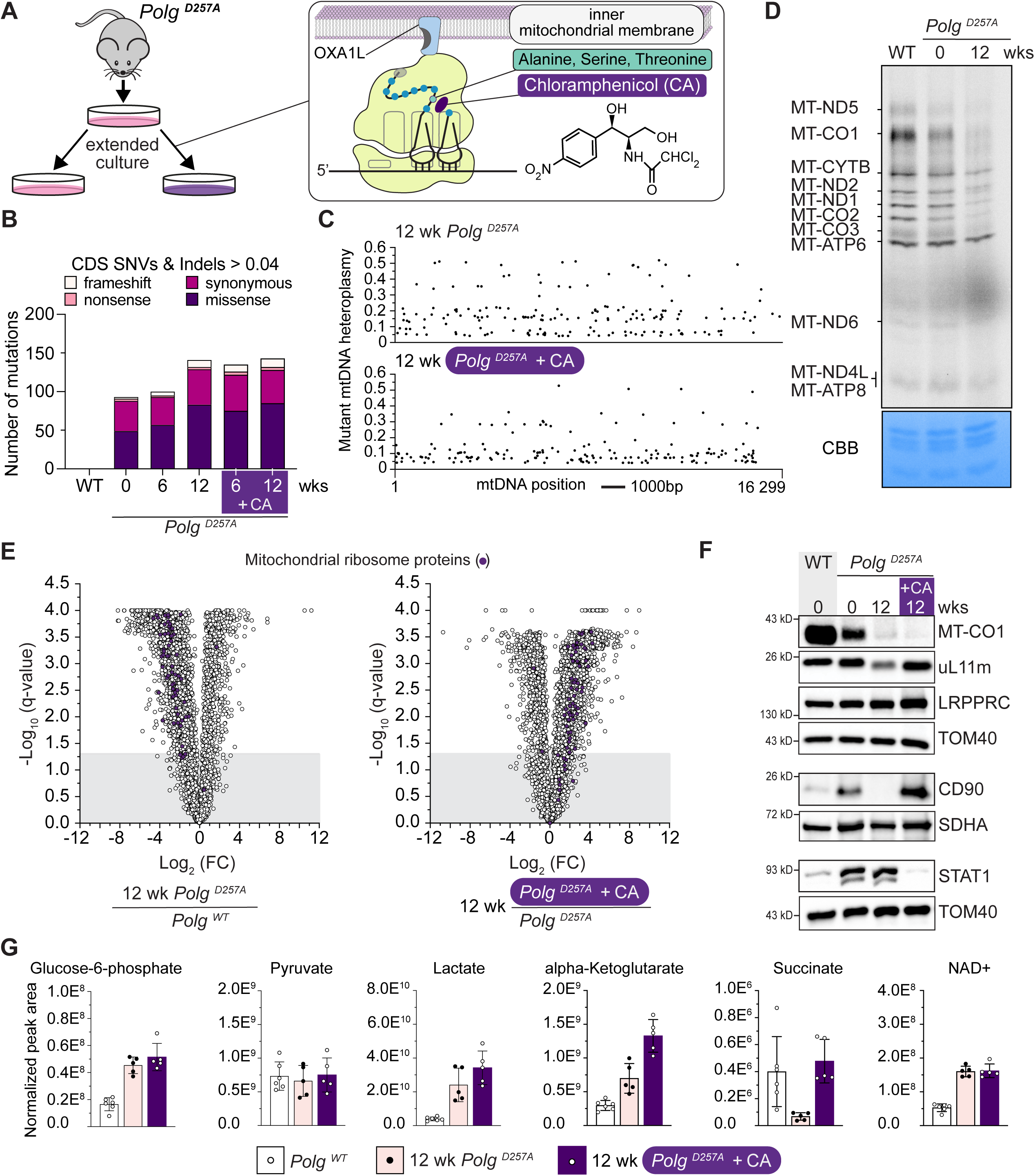
Inhibition of translation elongation on mitochondrial ribosomes reverses molecular phenotypes associated with the decline in cell fitness in *Polg ^D257A^* MEFs. (**A**) Schematic of the experimental design. CA, chloramphenicol. (**B**) Analysis of mtDNA mutations in coding sequence (CDS) from cultured cells that were greater than 0.04 in the cell population and (**C**) the distribution of all mutations along the mitochondrial genome. (**D**) A representative ^35^S-metabolic pulse labeling of mitochondrial protein synthesis. CBB, coomassie brilliant blue. (**E**) A volcano plot of protein abundance by quantitative label-free LC-MS/MS for the indicated genotypes comparisons (n=5). (**F**) Representative immunoblotting of whole cell lysates of MEFs with the indicated antibodies. (**G**) Metabolite levels determined from cells at 12 weeks of culture for *Polg ^D257A^* MEFs treated with and without chloramphenicol compared to *Polg ^WT^* MEFs. Each data point represents a separate cell sample. Bar graphs are mean +/- SD, n=5.

Long-term culturing of *Polg ^D257A^* MEFs was associated with a progressive increase in the mtDNA mutational burden and a decline in the synthesis rate of mitochondrial nascent chains and the abundance of mitochondrial ribosomes (Fig. 1B-F, fig. S2-S3 and table S1). Expression of this high mtDNA mutational burden profoundly altered nuclear gene expression (fig. S4, table S2) and the cellular proteome (Fig. 1E, table S3). Of note were significant shifts in the abundance of CD90, a stem cell marker integral to epidermal homeostasis and wound repair (*24*), and the innate immunity signaling factor STAT1 (Fig. 1E-F and table S3). Thus, our long-term culturing approach with *Polg ^D257A^* MEFs recapitulates hallmark molecular phenotypes associated with the decline in cell fitness that were previously established in cells and tissues derived from the mutator mouse model with age (*25–27*).

In contrast, chloramphenicol inhibition of mitochondrial protein synthesis significantly reversed many of these molecular signatures associated with the decline in cell fitness (Fig. 1E-F and fig. S5). MtDNA mutations still accumulated with chloramphenicol (Fig. 1B and fig. S2); whereas a single nucleotide mutation in the mouse mitochondrial rRNA *RNR2*, known to suppress the chloramphenicol molecular mode of action (*28*), was never detected in our analysis above a frequency of 0.01 (table S1). The data suggest that expression of mitochondrial mutations at the RNA level and the translation of these transcripts is a significant catalyst for the decline in cell fitness.

Mitochondrial dysfunction requires metabolic rewiring as part of compensatory feedback for cell survival (*27*, *29–32*). Classically, this mechanism is interpreted through the cellular demand for OXPHOS as seen with an increase in lactate (Fig. 1G and figS5). *Polg ^D257A^* cells are known to have altered expression levels of glycolytic enzymes (*25*) (fig S5), however, we posited that for chloramphenicol to reverse the molecular phenotypes associated with mtDNA mutations a distinct metabolic compensation would be required. To this end, chloramphenicol treatment of the *Polg ^D257A^* MEFs led to a robust fold increase in α-ketoglutarate levels (αKG) (Fig.1G), a cellular metabolite known to be critical for the pluripotency of stem cells and epigenetic regulation (*33*). Consistent with the mitochondrial translation dependent changes in αKG levels were FTO, an αKG-dependent mRNA demethylase that regulates cytosolic translation, and its substrate YAP1 (fig. S5). Recently, mitochondrial αKG metabolism was shown to regulate renewal of the intestinal stem cell niche (*34*). Maintenance of high αKG levels typically involves the catabolism of glucose and glutamine (*33*). The abundance of selected solute carriers was also regulated by chloramphenicol inhibition as part of the metabolic rewiring within the cell, including glutamine transporters (e.g., SLC38A family) at the plasma membrane (fig. S6). Remodeling of the *Polg ^D257A^* MEFs proteome is not driven strictly by transcriptional program (fig. S4), suggesting cytosolic translation and post-translational regulatory mechanisms in agreement with the significant shift in αKG levels. Thus, removing the molecular stress that arises from aberrations in mitochondrial protein synthesis allows the cell to counterbalance a severe bioenergetic defect in OXPHOS to maintain cell fitness, and in the process reveals an underappreciated complexity to organelle dysfunction and metabolism.

To further probe the capacity of mitochondrial protein synthesis errors to act as the specific trigger to these organelle and intracellular responses, we removed chloramphenicol from the media following 12 weeks for additional culturing (Fig. 2A). Within minutes of the chloramphenicol washout, the accumulated mtDNA mutations in the *Polg ^D257A^*MEFs (fig. S2) inhibited mitochondrial nascent chain synthesis despite wild type levels of the ribosomal protein uL11m (Fig. 2B and C). Persistence of the mitochondrial protein synthesis defect over the next three weeks directly affected the abundance of uL11m, CD90 and STAT1 (Fig. 2C). The subsequent reduction in uL11m was independent of mtDNA-encoded rRNA mutations that could impair subunit assembly (fig. S7). Instead, the data suggest that aberrations arising during translation elongation along mitochondrial mRNA transcripts induce the subsequent loss of mitochondrial ribosomal proteins over time in the *Polg ^D257A^* MEFs (Fig.1, 2, fig. S3 and table S2). Consistent with this interpretation is that impairments in the quality control of mitochondrial nascent chains are also known to activate mitochondrial ribosomal decay (*11*, *12*, *14*). Collectively, these data demonstrate that translation aberrations *per se* on mitochondrial ribosomes contribute to the impingement on cell fitness independently of OXPHOS.

**Fig. 2.**
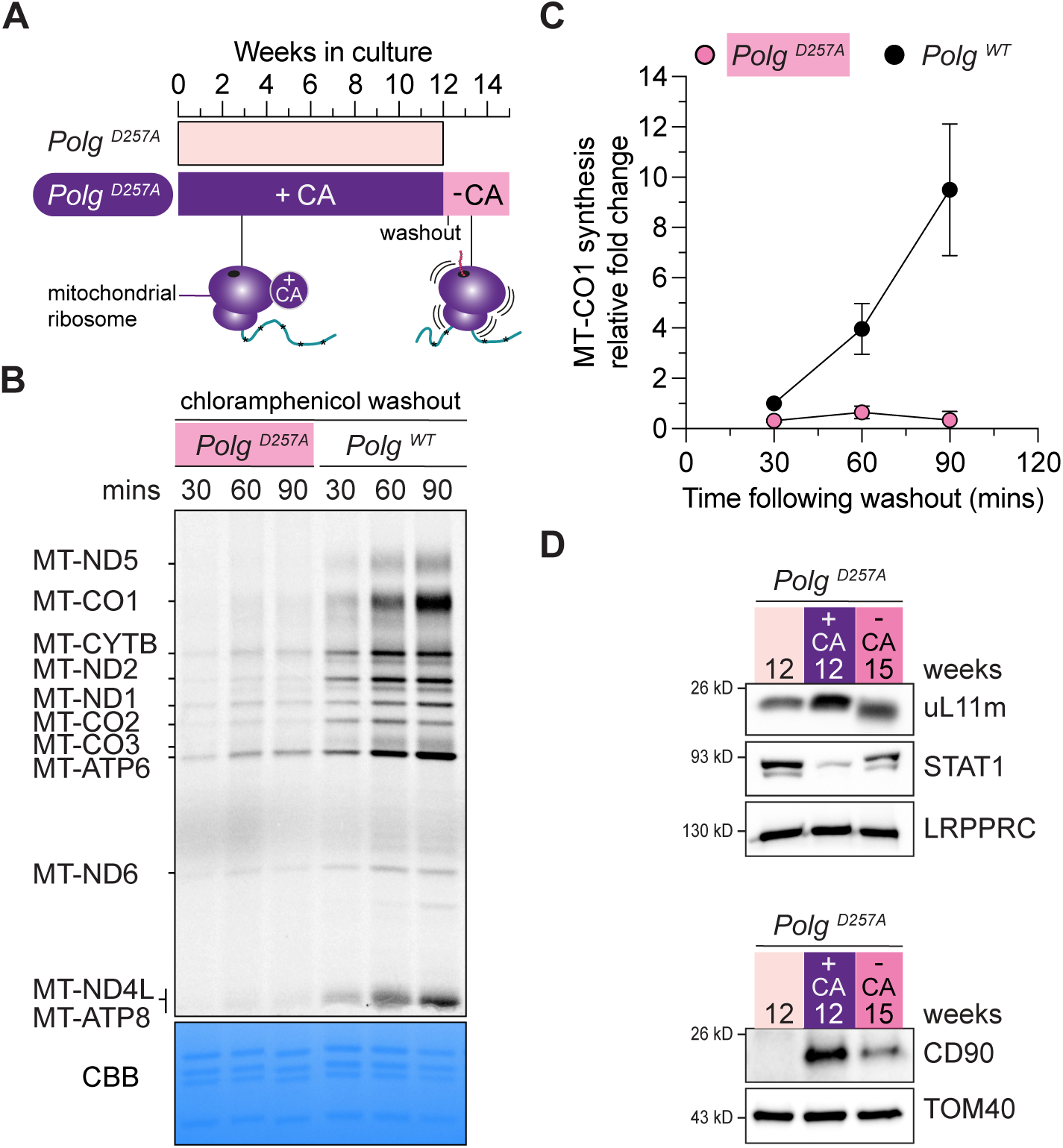
Translation of mitochondrial mRNAs encoding a high mutational burden impair protein synthesis and cell fitness. (**A**) Schematic of the experimental design. CA, chloramphenicol. (**B**) A representative ^35^S-metabolic pulse labeling of mitochondrial protein synthesis following the removal of chloramphenicol immediately from the media in MEFs with the indicated genotypes. In the *Polg ^D257A^* MEFs, chloramphenicol was removed from the media following 12 weeks of continuous culture with the antibiotic. In contrast, *Polg ^WT^* MEFs were treated with chloramphenicol for 24 hours before the antibiotic washout. CBB, coomassie brilliant blue. (**C**) Quantification of ^35^S-metabolic pulse labeling of mitochondrial protein synthesis following the removal of chloramphenicol in MEFs with the indicated genotypes. Data is a mean +/- SD. n=3. (**D**) Representative immunoblotting of whole cell lysates of MEFs following 12 weeks of extended culture and following the chloramphenicol washout 3 weeks later with the indicated antibodies.

At the steady state level, the majority of mtDNA in proliferating cell lines appears to be compacted into a tight nucleoid state regulated by TFAM that would be inaccessible to DNA replication and transcription factors (*35*). Although this genome architecture is dynamic, shifting towards open confirmations with mitochondrial dysfunction. Hence, the compaction state of the mitochondrial nucleoid could influence the frequency and types of mtDNA mutations expressed at the RNA level. TFAM expression in *Polg ^D257A^* MEFs was not affected with long-term culture or chloramphenicol inhibition of protein synthesis (fig. S8). All identified mtDNA mutations (> 0.01) were also detected at the RNA level and were significantly correlated in expression (fig. S8). Chloramphenicol had no effect on the RNA expression level of mtDNA encoded mutations (fig. S8).

In the *Polg ^D257A^* mouse, purifying selection prevents transmission of certain classes of deleterious mtDNA mutations from mother to offspring (*36*) via unknown and highly debated molecular mechanisms. Therefore, we asked whether aberrations in mitochondrial protein synthesis affected the segregation of mtDNA mutations. Distinguishing a genuine selective pressure against a mtDNA mutation from genetic drift, however, is a formidable challenge of mitochondrial genetics even for inherently deleterious variants (*37*). Since there is no homologous recombination of the mitochondrial genome (*38*), we focused on the segregation pattern of linked *in-cis* mtDNA mutations that had accumulated during 12 weeks of chloramphenicol treatment and the consequence following the washout as this would create an acute selective pressure. Over time, we observed altered segregation patterns of specific mtDNA haplotypes but not others with mitochondrial protein synthesis (fig. S9). We conservatively interpret these mtDNA segregation patterns as consistent with genetic drift.

Mitochondria play a central role in immunity (*39*, *40*). On the one hand mitochondria act as signaling hubs to mount antiviral responses; while on the other unique features arising from mitochondrial gene expression provide a ready source of damage-associated molecular patterns (DAMPs) (e.g., mtDNA and fMet peptides) that can be released following disruptions to the organelle integrity (*41*). A growing body of evidence points to the importance of mtDNA stress as a molecular trigger to potentiate immunometabolic dysfunction, dysregulating the cellular response to pathogen challenges (*25*, *41–45*). Thus, we asked what role mitochondrial protein synthesis plays in these innate immunity responses as the nexus step in gene expression.

A hallmark feature of the *Polg ^D257A^* mice is persistent type I interferon (IFN-I) activation due to heightened mtDNA instability, release and cGAS-STING activation (*25*, *45*). These interferon responses suppress activation of nuclear factor erythroid 2–related factor 2 (NRF2), leading to hyperinflammation that contributes to accelerated aging (*25*). NRF2 is a transcription factor that acts as a molecular rheostat to regulate cellular redox metabolism and control proinflammatory cytokine production (*46*). Extended culture of the *Polg ^D257A^*MEFs significantly affected the proteome of innate immunity pathways, including NRF2 (Fig. 3A-B and fig. S10). Chloramphenicol inhibition of mitochondrial protein synthesis in *Polg ^D257A^* MEFs robustly reversed these phenotypes back to wild type levels (Fig. 3A-B and fig. S10). Next, we explored the effect of chloramphenicol on bone marrow derived macrophages (BMDM) from old *Polg ^D257A^*mice (*25*), which have a higher mtDNA mutational burden than MEFs that continues to accumulate in culture despite a severely impaired growth rate (fig. S11). Chloramphenicol treatment reversed many of the IFN-I signaling phenotypes, including NRF2 suppression in BMDMs (fig.S12 and table S5). Thus, despite the biochemical inhibition of OXPHOS, chloramphenicol appears to drive a restoration of immunocompetence in *Polg ^D257A^* cells (Fig. 3A and fig. S11 and S12).

**Fig. 3.**
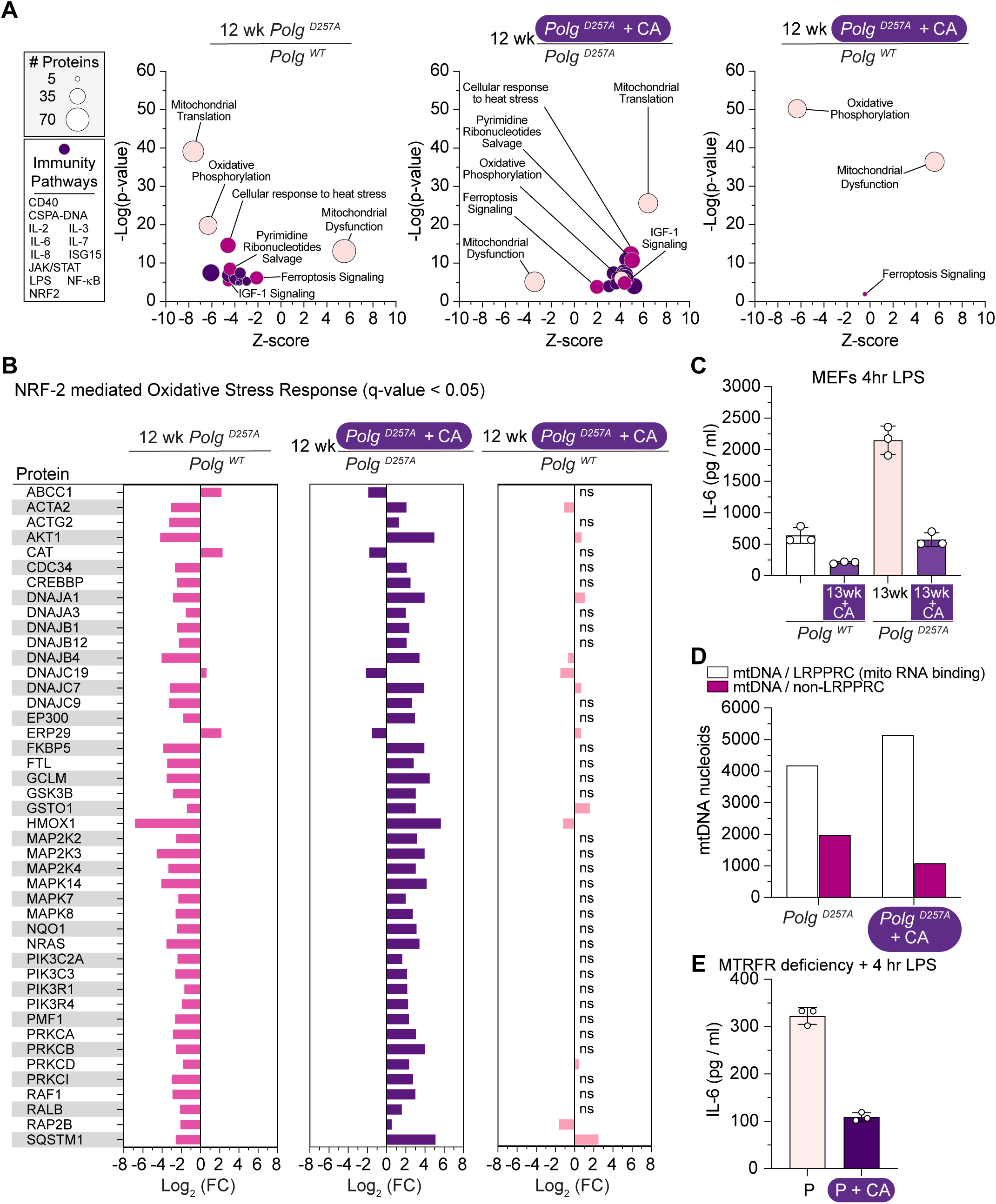
Aberrations during translation elongation on mitochondrial ribosomes trigger dysregulation of innate immune signaling. (**A**) Significantly enriched functional pathways (Ingenuity canonical pathways) of quantitative label-free LC-MS/MS data from MEFs with the indicated genotypes. Individual immunologically related pathways not labeled for sake of clarity: CD40; Cytosolic sensors of pathogen-associated DNA; IL-2; IL-3; IL-6; IL-7; IL-8; ISG15 antiviral; JAK/STAT, NRF2; NF-kappaB; LPS. CA, chloramphenicol. (**B**) Differential protein abundance in MEFs of the NRF-2 mediated oxidative stress response pathway determined from the Qiagen Ingenuity Pathway Analysis category of canonical pathways with the indicated genotypes and treatment conditions. Protein abundance was determined by label free LC:MS/MS with a q-value < 0.05. ns, non-significant. (**C**) IL-6 secretion into the media quantified by ELISA following 4 hrs of LPS challenge with MEFs with the indicated *Polg* alleles and treatment conditions. (**D**) Quantification of mtDNA puncta overlap with the mitochondrial mRNA binding protein LRPPRC in MEFs with *Polg^D257A^* and treatment conditions. A total of 6134 DNA puncta were quantified from untreated *Polg^D257A^* MEFs (n=23 cells) and 6201 DNA puncta from chloramphenicol treated *Polg^D257A^* MEFs (n=17 cells). (**E**) IL-6 secretion into the media quantified by ELISA following 4 hrs of LPS challenge from human MTRFR deficient fibroblasts with and without chloramphenicol treatment.

To functionally test this interpretation, we challenged the *Polg ^D257A^* MEFs with bacterial lipopolysaccharide (LPS) for 4 hours then measured the secretion of the pro-inflammatory cytokine interleukin 6 (IL-6). Expression of pathogenic *Polg* variants, including *Polg ^D257A^*, can generate excessive IL-6 following exposure to pathogen-associated molecular patterns (*25*, *45*). Consistently, chloramphenicol repressed the secretion of IL-6 to wild type levels in the *Polg ^D257A^* MEFs (Fig. 3C). The response correlated with an increase in the co-localization of mtDNA nucleoids with the mitochondrial RNA binding protein LRPPRC (Fig. 3D), suggesting less cytoplasmic mtDNA release which is a known activator for cGAS-STING (*42*). Since dysregulation of innate immunity appears to be a feature of mitochondrial disease (*47*), we asked whether the mechanism we established in *Polg ^D257A^* cells extends to inherited human pathogenic variants. We focused on a distinct step in mitochondrial nascent chain synthesis: ribosome quality control connected with translation termination on non-stop mRNAs. MTRFR is a key regulator of this process and pathogenic variants in the gene are associated with a severe impairment in mitochondrial protein synthesis and OXPHOS (*9*, *48*, *49*). Chloramphenicol treatment repressed IL-6 secretion upon LPS challenge (Fig. 3E). Together, these data reveal that aberrations in mitochondrial nascent chain synthesis *per se* can predispose a cell to abnormal inflammatory responses towards pathogens and not the OXPHOS dysfunction.

Our data suggest that mitochondrial protein synthesis acts as a control point to integrate organelle gene expression dysfunction into signals for the activation of major intracellular stress signaling pathways. Further we postulated that the homeostasis of the inner mitochondrial membrane is a critical component to this mechanism. To test this hypothesis, we inhibited the function of two AAA (ATPases associated with diverse cellular activities) complexes required for nascent chain quality control on the outer and inner membrane. ATAD1 regulates the quality control of misfolded nascent chains in the outer membrane (*50*, *51*); whereas the AFG3L2 complex controls the quality control of the 13 mtDNA encoded nascent chains destined for insertion into the inner membrane (*11–13*). Only AFG3L2 inhibition activated the integrated stress response (ISR) (Fig. 4A), a central intracellular stress signaling pathway that reprograms nuclear gene expression (*52*). A well-known ISR target is GDF-15, a potent biomarker of many disease states with underlying metabolic dysregulation (*53*). To explore this concept further, we turned to a powerful small molecule inducer of the ISR: actinonin (*14*, *30*, *54*). Actinonin is a formyl methionine mimetic, resembling the N-terminus of mitochondrial nascent chains, and a potent inducer of mitochondrial ribosome stalling (*5*, *11*, *14*). Actinonin triggered a robust secretion of GDF-15 into the media which could be completely blocked by co-treating with chloramphenicol (Fig. 4B). Collectively, these data show how the activation of another major intracellular stress signaling pathway arises specifically from aberrations in mitochondrial nascent chain synthesis and quality control.

**Fig. 4.**
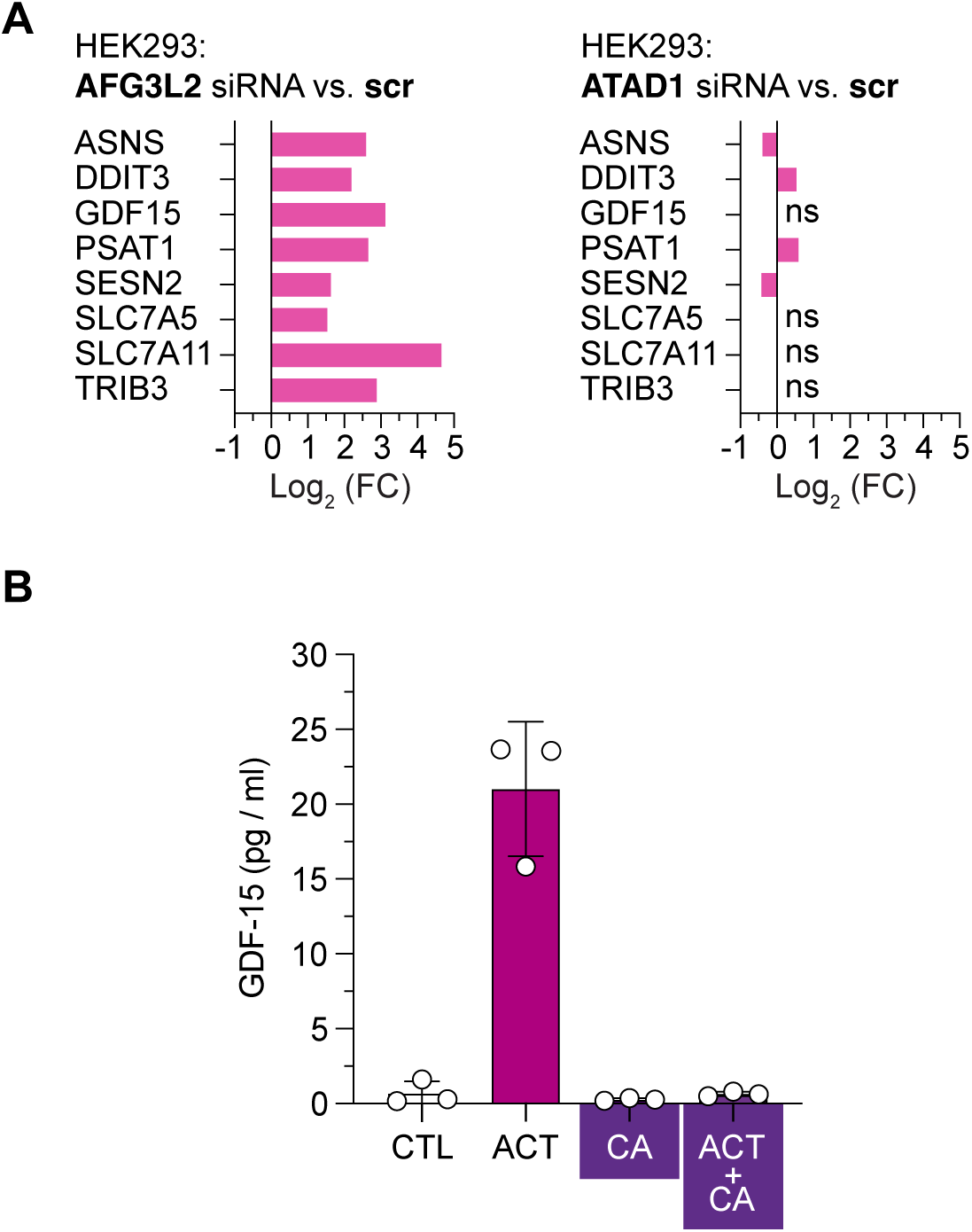
Aberrations during mitochondrial nascent chain quality control activate the integrated stress response and GDF-15 synthesis. (**A**) Differential mRNA expression of representative factors of ISR activation with mitochondrial dysfunction with a q-value < 0.05. HEK293 cells treated with small interfering RNA (siRNA) specific to the indicated essential subunits of mitochondrial AAA protein complexes anchored in the mitochondrial inner membrane (AFG3L2) or outer membrane (ATAD1) (**B**) GDF-15 secretion into the media quantified by ELISA following 24 hrs of actinonin (ACT) treatment, chloramphenicol (CA) or both.

Several important principles emerge from our findings. First, a bioenergetic defect in OXPHOS alone cannot account for all of the cellular phenotypes arising from mitochondrial dysfunction. Second, at the molecular level, the complete inhibition of nascent chain synthesis is fundamentally distinct on organelle function from errors arising during translation elongation and termination on mitochondrial ribosomes. The inherent ability of a given cell type to generate and resolve these errors is critical to homeostasis. Thus, understanding the fidelity of mitochondrial protein synthesis and how the co-translational insertion of mitochondrial nascent chains into the inner membrane can impinge on organelle function is paramount. Third, defects in mitochondrial protein synthesis can trigger the secretion of a myriad of cytokines with different modes of action. Elucidating the mechanisms and importance of this cell-to-cell communication is integral for deciphering the molecular pathogenesis of mitochondrial dysfunction. Lastly, there is a surprising complexity to how OXPHOS inhibition can rewire metabolism, exerting direct effects on the epigenome and translation control on cytoplasmic ribosomes that are linked to cell fitness. Together, our data demonstrate how defects in mitochondrial protein synthesis *per se* acts as a major molecular contributor to the decline in cell fitness associated with mitochondrial pathologies and to the aging process itself.

## METHODS

### Biological samples and cell culturing

Mouse embryonic fibroblasts were derived from E13.5 embryos generated by a single intercross of *Polg ^WT/D257A^* mice on a C57BL/6NCrl nuclear background (*1*) to minimize the mtDNA mutational burden (*21*). MEFs were cultured in Dulbecco’s modified Eagle’s medium (DMEM) (Merck) with high glucose supplemented with 10% fetal bovine serum (FBS), 1× glutamax, and uridine (50 μg/ml; Sigma-Aldrich, # U3003) at 37°C and 5% CO2. All cells tested negative for mycoplasma infection (PromoKine). MEFs were subsequently immortalized by retroviral transduction of E7 and hTERT (*55*). Primary bone marrow derived macrophages (BMDMs) were generated from femur and tibia bones of *Polg^WT^* and *Polg^D257A^* on a C57BL/6J nuclear background (*2*) and cultured on petri dishes in DMEM containing 10% FBS plus 30% L929 culture media for 7 days. L929 cells were obtained from ATCC and maintained in DMEM (D5796, Sigma) supplemented with 10% FBS (97068-085, VWR). For immortalization, BMDMs were infected with J2 recombinant retrovirus (*56*). Immortalized macrophages were maintained in culture for 2-3 months and were slowly weaned off L929 supernatant until they were growing in the absence of conditioned medium (*57*) in DMEM with 10% FBS, 1× glutamax, and uridine (50 μg/ml) at 37°C and 5% CO2 (*25*). Chloramphenicol (124 mM) (Merck) was added to the culture media every 3 days.

Two isogenic human myoblasts cell lines homoplasmic for the m.8344A>G or its matched wild type mtDNA and m.3243A>G or its matched wild type mtDNA were previously generated (*58*). Myoblasts were cultured in Skeletal Muscle Cell Growth Medium (Sigma-Aldrich, #151-500) supplemented with uridine 50 μg/ml at 37°C and 5% CO_2_. The HEK293 cell line was cultured in DMEM with 10% FBS, 1× glutamax, and uridine (50 μg/ml) at 37°C and 5% CO2. Stealth siRNAs against *AFG3L2* (HSS116886), *ATAD1* (HSS131410) and scrambled sequences were obtained from Life Technologies and transfected on day 1 and 3 of culture with Lipofectamine^TM^ RNAiMAX (Thermo Fisher) and collected on day 5 for analysis. All siRNA knockdowns were evaluated by immunoblotting using specific antibodies.

### Antibodies

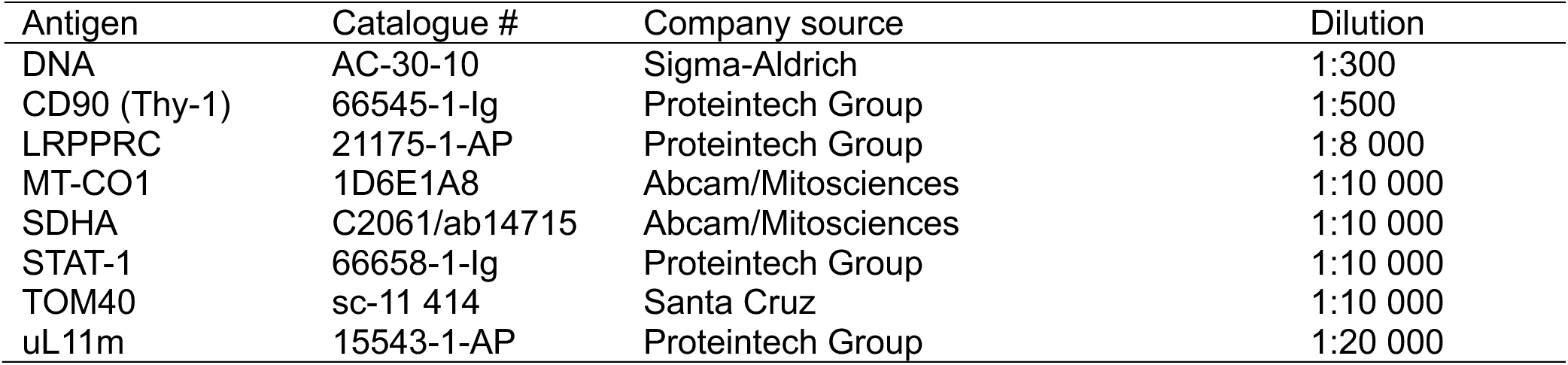

| Antigen | Catalogue # | Company source | Dilution |
| --- | --- | --- | --- |
| DNA | AC-30-10 | Sigma-Aldrich | 1:300 |
| CD90 (Thy-1) | 66545-1-Ig | Proteintech Group | 1:500 |
| LRPPRC | 21175-1-AP | Proteintech Group | 1:8 000 |
| MT-CO1 | 1D6E1A8 | Abcam/Mitosciences | 1:10 000 |
| SDHA | C2061/ab14715 | Abcam/Mitosciences | 1:10 000 |
| STAT-1 | 66658-1-Ig | Proteintech Group | 1:10 000 |
| TOM40 | sc-11 414 | Santa Cruz | 1:10 000 |
| uL11m | 15543-1-AP | Proteintech Group | 1:20 000 |

### Immunoblotting

Cells were lysed in phosphate buffered saline, 1% dodecyl-maltoside (DDM), 1 mM phenylmethylsulfonyl fluoride (PMSF) and complete protease inhibitor (Thermo Fisher). Protein concentrations were measured with the Bradford assay (BioRad). Equal amounts of proteins were separated by tris-glycine SDS-PAGE and transferred to a nitrocellulose membrane by semi-dry transfer. Membranes were blocked at room temperature for 1 hour in TBST with 1% milk, followed by incubation with primary antibodies overnight at 4°C in 5% BSA/TBST. Signals were developed on the following day with secondary anti-mouse or anti-rabbit HRP conjugates (Jackson ImmunoResearch) using enhanced chemiluminescence with the c300 Chemiluminescent imaging system (Azure Biosystems). Representative data of independent experiments were cropped in Adobe Illustrator after linear adjustments.

### Immunostaining

Cells plated on coverslips were washed with phosphate-buffered saline (PBS) three times before fixation with 4% paraformaldehyde (PFA) in PBS for 15 minutes at room temperature followed by 3X washing in PBS, permeabilization with 100% methanol followed by 3X washing with PBS. Coverslips were then incubated with 5% bovine serum albumin (BSA) in PBS containing 0.1% Tween®20 (Sigma-Aldrich) for 1 hour at room temperature. Primary antibodies were diluted in 5% BSA/PBST and incubated on coverslips overnight at 4°C. The following day, coverslips were washed three times with PBST. Secondary anti-mouse or anti-rabbit fluorescently labeled antibodies in 5% BSA/PBST were incubated for 1 hour at room temperature. Coverslips were washed three final times with PBST. Hoechst 33342 (1:2000, Invitrogen) was included in the first wash to stain nuclear DNA. Coverslips were mounted with ProLong™ Glass Antifade Mountant (Thermo Fisher) on microscopy slides and incubated overnight at 4°C before imaging.

### Microscopy

Slides were imaged using an inverted microscope (Olympus FV4000) at room temperature (RT) with a UPLXAPO 60×/1.42 oil immersion objective (WD 0.15 mm) and OBIS laser (640 nm). Images were acquired using a resonant scanner at 1024 × 1024 pixels (roundtrip) with cellSens FV imaging software and exported into ImageJ using the Fiji plugin for brightness and contrast adjustments.

For live-cell imaging for growth rate analysis, cells were seeded in 6-well plates at a density of 6 × 10⁴ cells per well on day 0. The plates were transferred to the Cell-IQ imager, and phase-contrast images were taken from each plate to obtain 16 frames per plate every 2 hours, using a Nikon objective, during 5 days for BMDMs and 3 days for MEFs. The images were analyzed with Cellpose software using the Cyto3 model to count the total number of cells per well from all 16 frames.

To determine growth rates, cells were seeded to 6 well plates with 6x10^4^ cells/well. Cell proliferation was determined using the Cell-IQ imager. Images were analyzed with the Cellpose software using the Cyto3 model to count total number of cells per well obtained.

### MtDNA localization and quantification

Confocal microscopy images taken following immunostaining with anti-LRPPRC, anti-DNA and DAPI were analyzed with SpotMAX integrated to Cell-ACDC software (*59*) for the counting mtDNA nucleoids and organelle localization. After creating data structure and launching data prep module with default settings, cells were segmented on the LRPPRC channel using cellpose_V3 model and cyto3 model setting diameter=250 and Gaussian filter=0.5 in the pre-processing. In the first step, mtDNA nucleoids were detected in reference to the LRPPRC signal by segmenting the channel and counting DNA puncta for overlap or not. Second, the DAPI channel was segmented to exclude any DNA puncta from the collection that were clearly of nuclear origin.

### Radioisotope labeling of mitochondrial translation

Cultured cells were pre-treated with anisomycin (100 μg/ml) for 5 minutes to inhibit cytoplasmic translation then pulsed with 200 μCi/ml ^35^S Met-Cys (EasyTag-Perkin Elmer) for 30 minutes. Equal amounts of sample protein determined via the Bradford assay were treated with Benzonase (Thermo Fisher) on ice for 20 minutes and resuspended in 1x translation loading buffer (186 mM Tris-Cl pH6.7, 15% glycerol, 2% SDS, 0.5 mg/ml bromophenol blue, 6% beta-mercaptoethanol). A 12-20% gradient tris-glycine SDS-PAGE was used to separate labelled proteins and dried for exposure with a phosphorimager screen and scanned with the Azure Biosystems imaging platform for quantification. Gels were rehydrated in water and stained Coomassie-blue to verify sample loading.

### Proteomics analysis

Cultured cell samples from MEFs and BMDM were subjected to quantitative label-free proteomics analysis. Proteins were precipitated on amine beads as previously described (*60*). The precipitated proteins on beads were dissolved in 50 mM ammonium bicarbonate, reduced, alkylated and digested with trypsin (1:50 enzyme:protein ratio; Promega, United States) at 37°C overnight. The resulting peptide mixture was purified by STAGE-TIP method using a C18 resin disk (3M Empore) before the samples were analyzed by a nanoLC-MS/MS using nanoElute coupled to timsTOF PRO2 (Bruker) with 60-minute separation gradient and 25 cm Aurora C18 column. The times TOF PRO2 was operated in DDA (MEFs) or DIA (BMDM) mode. MS raw files from MEFs were submitted to MaxQuant software version 2.4.7.0 for protein identification and label-free quantification. Carbamidomethyl (C) was set as a fixed modification and acetyl (protein N-term), carbamyl (N-term) and oxidation (M) were set as variable modifications. First search peptide tolerance of 20 ppm and main search error 10 ppm were used. Trypsin without proline restriction enzyme option was used, with two allowed miscleavages. The minimal unique + razor peptides number was set to 1, and the allowed FDR was 0.01 (1%) for peptide and protein identification. Label-free quantitation was employed with default settings. UniProt database with ‘mouse’ entries was used for the database searches. MS raw files from BMDM were submitted to DIA-NN (version 1.9.2) for protein identification and label-free quantification using the library-free function. The UniProt mouse database was used to generate library *in silico* from a FASTA file. Carbamidomethyl (C) was set as a fixed modification. Trypsin without proline restriction enzyme option was used, with one allowed miscleavage and peptide length range was set to 7-30 amino acids. The mass accuracy was set to 15ppm and precursor false discovery rate (FDR) allowed was 0.01 (1%). LC-MS/MS data quality evaluation and statistical analysis was done using software Perseus ver 1.6.15.0 and pathway analysis was done using Ingenuity Pathway Analysis (Qiagen).

### Metabolomics analysis

For extraction of intracellular metabolites, cells were washed in ice-cold PBS followed by extraction with ice-cold extraction buffer containing 80% acetonitrile in distilled water. Samples were centrifuged to collect the supernatant and stored at −80 °C until analysis. Briefly, 5 µL of sample was injected into a Thermo Dionex UltiMate 3000 HPLC system (Thermo Scientific) coupled with a Q Exactive Plus Orbitrap mass spectrometer (Thermo Fisher Scientific). Samples were run in randomized order with quality control samples and blanks inserted every tenth injection. HPLC was equipped with a hydrophilic ZIC-pHILIC column (150 × 2.1 mm, 5 μm) and a ZIC-pHILIC guard column (20 × 2.1 mm, 5 μm, Merck Sequant) for metabolite separation. Chromatographic separation was achieved using a gradient program with mobile phases comprising 200 mmol/L ammonium bicarbonate (pH 9.3, adjusted with 25% ammonium hydroxide), 100% acetonitrile, and deionized water. A linear solvent gradient was applied, decreasing the organic solvent from 80% to 35% over 16 minutes, at a flow rate of 0.15 mL/min and a column oven temperature of 45°C. The ammonium bicarbonate solution was kept at 10% throughout the run, resulting in a constant concentration of 20 mmol/L. A Q Exactive Plus Orbitrap mass spectrometer (Thermo Fisher Scientific) equipped with electrospray ionization was coupled to the HPLC system for metabolite profiling and identification. For metabolite profiling, polarity switching mode was employed with a resolution (RES) of 70,000 (at 200 m/z) to detect both positive (3,400 V) and negative (3,000 V) ions across a mass range of 75-1,000 m/z. Settings included a sheath gas of 28 arbitrary units (AU) and auxiliary gas of 8 AU; vaporizer temperature of 280°C; and ion transfer tube temperature of 300°C. The instrument was controlled using Xcalibur software version 4.1.31.9 (Thermo Scientific). Metabolite peaks were validated by comparison to commercial standards (Sigma-Aldrich). Data quality was assessed throughout the run using an in-house quality control cell line, processed as previously described (*61*). Raw data were analyzed using Skyline (*62*). Compounds were identified by matching the mass and retention time of detected peaks to an in-house library generated from metabolite standards (mass tolerance of 5 ppm and retention time tolerance of 2 minutes).

### Mitochondrial DNA sequencing and analysis

Mitochondria were isolated from cell pellets using the Qproteome Mitochondria Isolation Kit (Qiagen) following the manufacturers’ methods. DNA was isolated using QIAamp DNA Kit (Qiagen). Three amplicons covering the entire mitochondrial genome were generated by PCR using the following primer pairs.

Amplicon 1:

GLB57 5’- TTCTTCTAAAATTAGGTAGTTACGGAATAATTCGCATCTCCATTATTCTA

GLB53 5’- TTGTTAATGTTTATTGCGTAATAGAGTATGATTAGAGTTTTGGTTCACGG

Amplicon 2:

GLB55 5’- TAATTCGAGCAGAATTAGGTCAACCAGGTGCACTTTTAGGAGATGACCAA

GLB56 5’- CATAGGGAGAGAAGGATGAAGGGGTATGCTATATATTTTGTTAGTGGGTC

Amplicon 3:

GLB52 5’- GTTAATGTAGCTTAATAACAAAGCAAAGCACTGAAAATGCTTAGATGGAT.

GLB80 5’- TTCTACTATTGATGATGCTAGGAGAAGGAGAAATGATGGTGGTAGGAGTC.

For short read Illumina sequencing, mtDNA amplicons were fragmented for library construction followed by Illumina NovaSeq 2 X 150 bp deep sequencing for 10M paired reads. Analysis of deep sequencing reads for mitochondrial DNA variants was performed according to standard approaches, using only R1 reads to maintain high quality that enables identification of rare variants (*63*). Raw reads were quality and adapter trimmed using flexbar v.3.5.0 (-qt 28 -m 50 -ao 10 -at ANY -ae 0.1) and then mapped on to the *Mus musculus* C57BL/6J reference mitochondrial genome (NC_005089.1) (bwa mem v.0.7.17 with options -T 19 -B 3 -L 5,4). Trimmed reads were also mapped on a ‘split-genome’ generated by cutting the genome in half and joining the ends to account for the circular nature of the mtDNA. Mapped reads were sorted with ‘sort’ and unmapped reads were filtered out using ‘view’ from samtools v.1.15. InDel qualities were re-calculated with lofreq indelqual. Variants were called and filtered with lofreq call-parallel (-N -B -q 30 -Q 30) and lofreq filter (--no-defaults -- snvqual-thresh 70 --indelqual-thresh 70 --sb-incl-indels -B 60) of lofreq v.2.1.4. Variants were further filtered with SnpSift (DP*AF >= 15, DP4[2] >= 3, DP4[3] >=3) before annotating with SnpEff v.5.1 (-no-downstream -no-upstream). A final filtering was applied for strand bias (<1000) and allele frequency (0.001>af>0.7) with custom python3 scripts. Depth of coverage for each position were extracted using bedtools v.2.30. Depth values from the first and last 200nt’s were taken from the split-genome-mapped reads. Various other data handling was done with samtools and python3.

For short read Illumina sequencing, mtDNA amplicons were fragmented for library construction followed by Illumina NovaSeq 2 X 150 bp deep sequencing for 10M paired reads. Analysis of deep sequencing reads for mitochondrial DNA variants was performed according to standard approaches (*63*). Briefly, only R1 reads were used to maintain high quality. Raw reads were quality and adapter trimmed using flexbar v.3.5.0 (-qt 28 -m 50 -ao 10 -at ANY -ae 0.1). Trimmed reads were then mapped on to the *Mus musculus* mitochondrial genome (NC_005089.1), as well as a “split-genome” made from this genome by cutting the genome in half and merging the start-end positions together (bwa mem v.0.7.17 with options -T 19 -B 3 -L 5,4). Read depth artificially drops at both ends of the linear reference sequence of the circular mitochondrial genome; thus, a “split-genome” approach was used accordingly (*63*). Mapped reads were sorted with ‘sort’ and unmapped reads were filtered out using ‘view’ from samtools v.1.15. InDel qualities were re-calculated with lofreq indelqual before calling variants with lofreq call-parallel with lofreq v.2.1.4 (-N -B -q 30 -Q 30). Variants were filtered with lofreq filter (--no-defaults --snvqual-thresh 70 --indelqual-thresh 70 --sb-incl-indels -B 60). Variants from reference and split-reference genome were merged such that the variants from the first and last 200nt’s would come from the split-genome approach. Single Nucleotide Variants (SNVs) and InDels were separated using lofreq filter and both were filtered again with SnpSift from SnpEff v.5.1 (DP*AF >= 15, DP4[2] >= 3, DP4[3] >=3). Finally, variants were annotated using SnpEff v.5.1 (-no-downstream -no-upstream). A final filtering was applied for strand bias (<1000) and allele frequency (0.001>af>0.7) with custom python3 scripts. Depth of coverage for each position were extracted using bedtools v.2.30. Depth values from the first and last 200nt’s were taken from the split-genome-mapped reads. Various other data handling was done with samtools and python3.

For long read sequencing of mtDNA amplicons, PacBio SMRTbell amplicon libraries for PacBio Sequel were constructed using SMRTbell prep kit 3.0 (PacBio, Menlo Park, CA, USA) using the manufacturer recommended protocol. Primer annealing, polymerase binding, and complex cleanup was performed using the Sequel binding kit 3.1 & cleanup beads (PacBio) and loaded onto PacBio Sequel IIe. Sequencing was performed on PacBio Sequel IIe SMRT cell. Sequel IIe sequencing generated subread files, as well HiFi reads via on-board CCS analysis. Demultiplexing was performed using PacBio lima.

PacBio HiFi reads were analyzed using the Python3 package pysam v0.22.1. Variant sites identified in the Illumina dataset were queried using the pileup method with a minimum base quality threshold of 40 applied to ensure high-confidence variant detection (*64*). The approach enabled direct comparison of heteroplasmy detection between short-read and long-read sequencing platforms using Python3 with the scipy.stats.pearson function. For haplotype analysis, a stricter per-read filter was applied, retaining only reads with base quality >40 at all queried heteroplasmic positions. This criterion ensured that haplotype-level analyses were based solely on reads with high-confidence base calls across all relevant loci. To rule out template switching during PCR of the mtDNA amplicons as a technical artefact, we generated an amplicon from total heart DNA isolated from a heteroplasmic mouse model with the NZB and BALB/c mtDNA haplotypes (*65*). Since there is no homologous recombination of mtDNA (*38*), this DNA sample provided a technical control to determine PCR artifacts. The resulting error estimate was then applied as a threshold for interpreting linkage in the *Polg ^D57A^* mtDNA amplicons.

### RNA sequencing

For each MEF sample (n=1), total cell RNA was isolated from a single 150 mm plate of cultured cells with the Reliaprep RNA Miniprep kit (Promega) followed by deoxyribonuclease I (NEB) treatment. Sequencing libraries were constructed accordingly (*9*). Illumina reads were preprocessed with Trimmomatic v.0.39 (SE mode; parameters: MINLEN:30, HEADCROP:1). For mitochondrial transcript analysis, reads were aligned to the Mus musculus mitochondrial reference genome (NC_005089.1) using BWA MEM v.0.7.18 to retain polycistronic transcripts. Mapped reads were processed with Samtools mpileup v.1.21 to generate nucleotide-resolution coverage profiles across the mitochondrial genome. Nuclear gene expression was quantified by aligning reads to the mm39 reference transcriptome using the pseudoaligner Salmon v.1.9.0. Differential expression analysis was performed using python3 packages pytximport and pydeseq2.

For human RNA sequencing analysis, each sample (n=1) consisted of total cell RNA isolated from a single 100 mm plate using the Reliaprep RNA Miniprep kit (Promega) followed by deoxyribonuclease I (NEB) treatment. DNA library preparation was performed using NEBNext Ultra DNA Library Prep kit following the manufacturer’s instructions (Illumina, San Diego, CA, USA). The libraries were pooled equimolar and loaded with a 30% spike-in of PhiX Sequencing Control V3 on a S4 flowcell on the Illumina NovaSeq 6000 instrument according to manufacturer’s instructions. The samples were sequenced using a 2x150 paired end (PE) configuration. Image analysis and base calling were conducted by the NovaSeq Control Software v1.7 on the NovaSeq instrument. Raw sequence data (.bcl files) generated from Illumina NovaSeq was converted into FASTQ files and de-multiplexed using Illumina bcl2fastq program version 2.20. One mismatch was allowed for index sequence identification.

### Enzyme-linked immunosorbent assay (ELISA)

To measure IL-6 secretion upon lipopolysaccharide (LPS) challenge, 8x10^4^ MEFs or human fibroblasts were plated per well in a 12 well plate in 1ml media. The following day, the media was changed then cells were incubated with LPS (1ug/ml) (Invivogen, LPS-B5 Ultrapure) for 4 hours. Cell supernatants were centrifuged to remove cell debris and detached cells then analyzed using the ThermoFisher Scientific mouse IL-6 (Catalogue # BMS603-2) or human (Catalogue # EH2IL6) ELISA kits according to the manufacturer’s instructions. To measure GDF-15 secretion, cell culture supernatants were analyzed using the R&D systems Mouse and Rat GDF15 ELISA kit (catalogue # MGD150) according to the manufacture’s instruction. To induce GDF15 secretion, cells were treated with 150 μM actinonin a small molecule inhibitor of AFG3L2 and mitochondrial co-translational nascent chain degradation (*11*) that is known to induce the integrated stress response and GDF15 mRNA expression (*30*).

## Acknowledgements

The authors wish to acknowledge the support of the life science research infrastructures at the Institute of Biotechnology, HiLIFE, University of Helsinki: Viikki DNA Sequencing and Genomics, and Light Microscopy Unit (LMU). Furthermore, the mass spectrometry-based proteomic analyses were performed by the Proteomics Core Facility, Department of Immunology, University of Oslo/Oslo University Hospital, which is supported by the Core Facilities program of the South-Eastern Norway Regional Health Authority. This core facility is also a member of the National Network of Advanced Proteomics Infrastructure (NAPI).

## Author contributions

Conceptualization: GLB; KYN; AEE; BJB.

Data curation: GLB; EB; AA; LK; SN; AKN; TAN; BJB.

Formal analysis: GLB; EB; AA; LK; SN; AKN; TAN; BJB.

Investigation: GLB; KYN; EB; AEE; LK; SN; CR; TVT; TAN.

Methodology: GLB; JJV; JBS; APW.

Project administration: GLB; KYN; AEE; BJB.

Resources: SOK; JBS; JJV; APW.

Software: GLB; EB; AA; JBS.

Supervision: GLB; AEE; BJB

Validation: GLB; KYN; LK; BJB.

Visualization: GLB; BJB

Writing-original draft: GLB; BJB.

Writing-review and editing: GLB; KYN; EB; AA; AEE; LK; CR; TVT; SOK; TAN; JBS; APW; BJB

## SUPPLEMENTAL FIGURE LEGENDS

**fig. S1:**
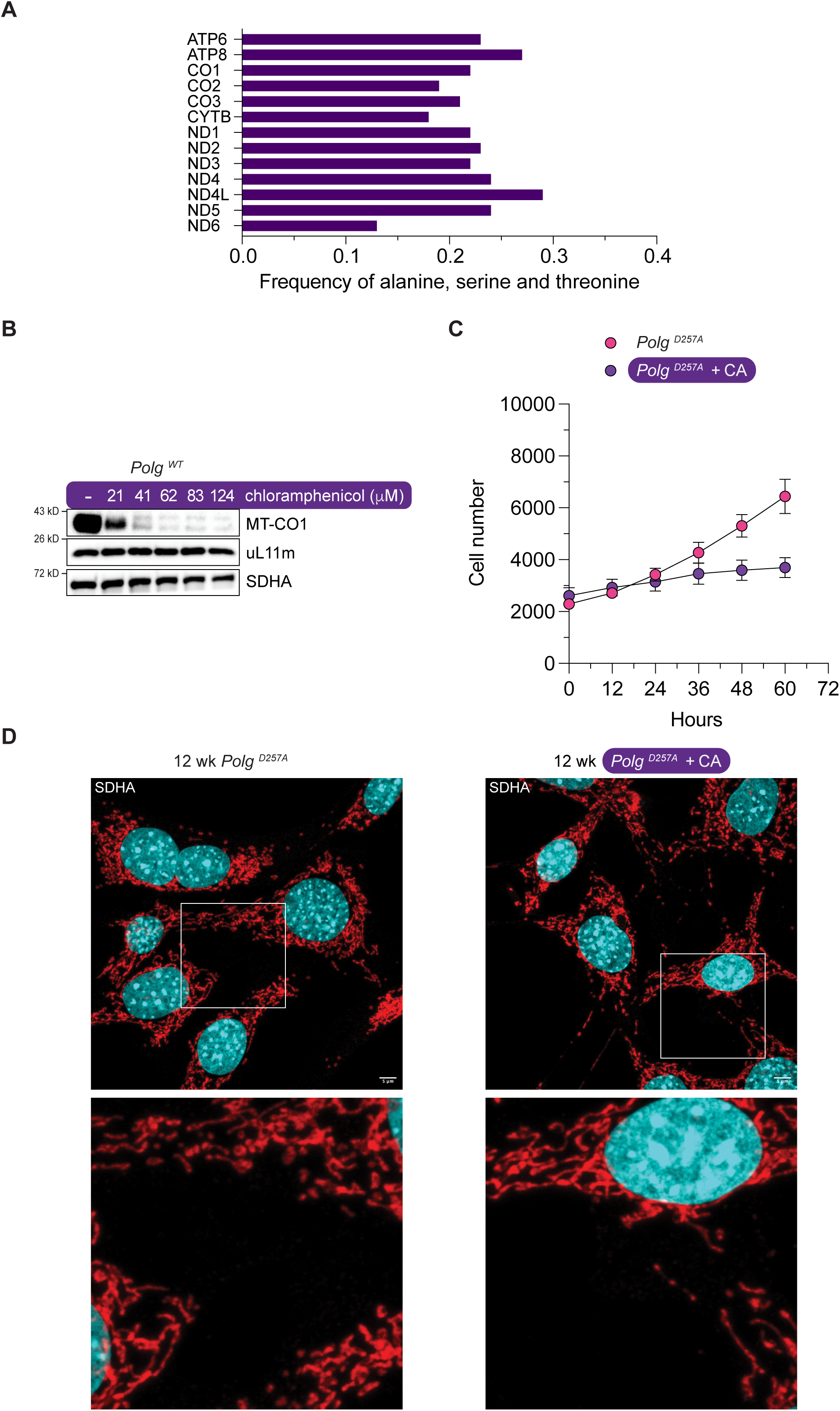
Chloramphenicol inhibition of mitochondrial translation in *Polg ^D257A^* mouse embryonic fibroblasts (MEFs). (**A**) Frequency of alanine, serine and threonine residues in the 13 protein coding genes of the mouse mtDNA. For chloramphenicol inhibition of translation elongation, one of three amino acids is required in the −1 position of the nascent polypeptide chain. (**B**) A chloramphenicol dose-response curve of MEFs with immunoblotting against the indicated proteins after 5 days of treatment. MT-CO1 is encoded in mtDNA. uL11m is a nuclear encoded mitochondrial ribosomal protein of the large subunit. SDHA is a nuclear encoded subunit of complex II of the oxidative phosphorylation complexes. (**C**) Growth curve of 12-week-old *Polg ^D257A^* MEFs with and without chloramphenicol (CA). Cells were seeded at equal densities and imaged over 60 hours by live-cell imaging. (**D**) Representative immunostaining images showing mitochondrial morphology of 12-week-old *Polg ^D257A^* MEFs with and without chloramphenicol.

**Fig. S2:**
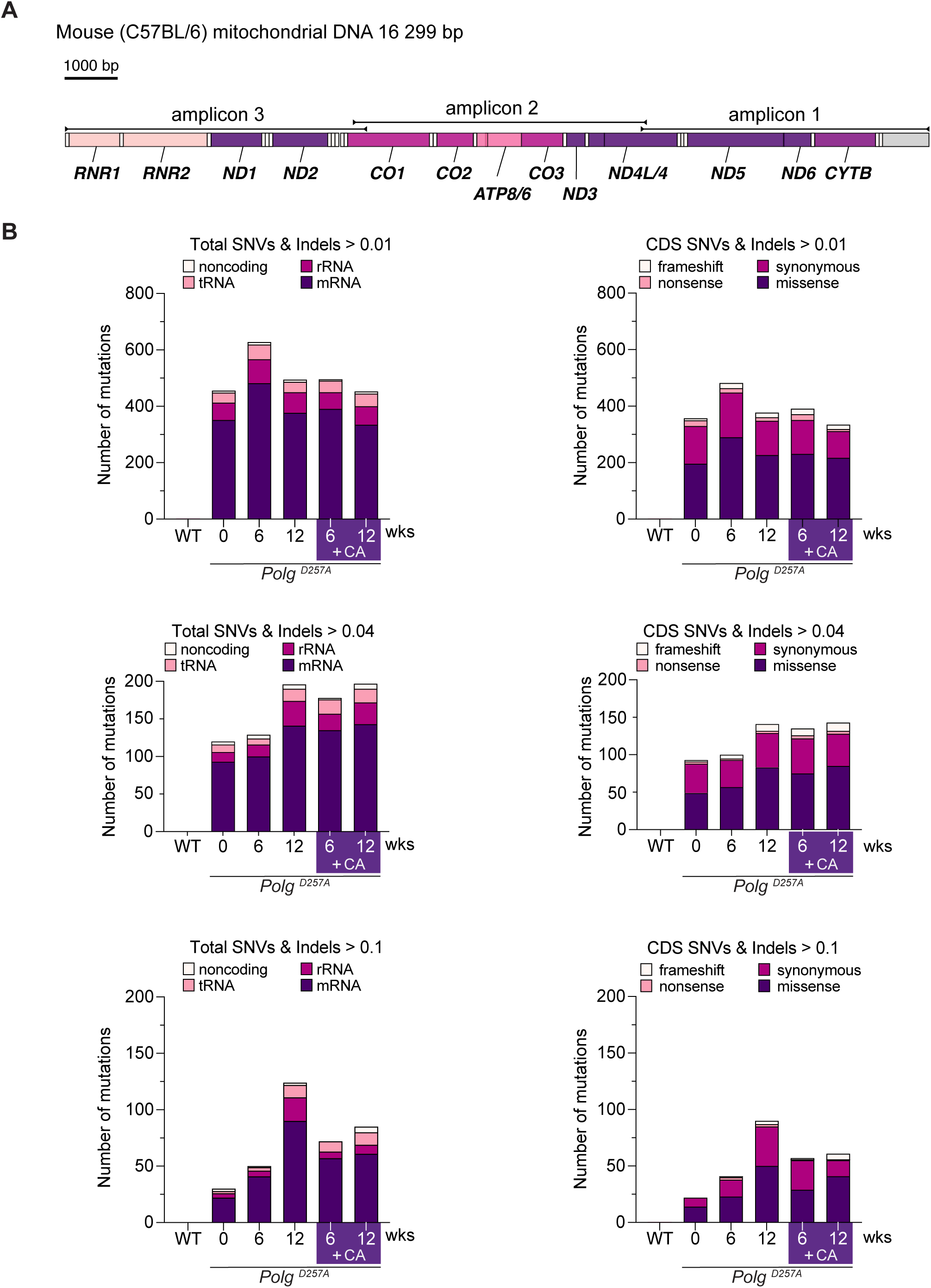
Mitochondrial DNA mutations accumulate in *Polg ^D257A^* MEFs with extended culture. (**A**) Schematic of the mouse mtDNA with location of amplicons indicated that were generated for deep sequencing mutation detection. (**B**) Mutation analysis of mtDNA isolated from *Polg ^D257A^* MEFs following extended culture occurring greater than 0.01, 0.04 and 0.1.

**Fig. S3:**
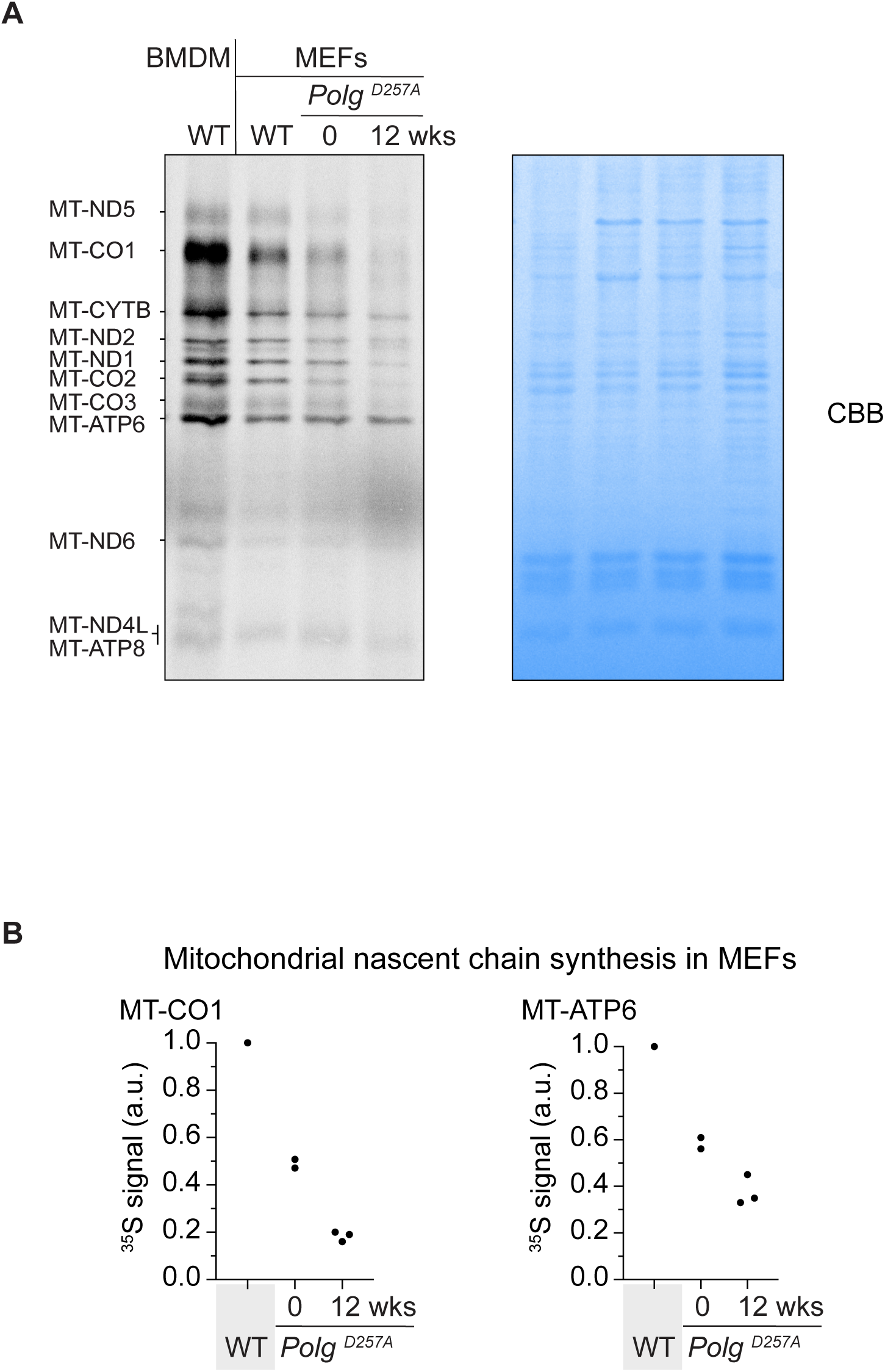
Mitochondrial DNA mutational load accumulated with long-term culturing of *Polg ^D257A^*MEFs progressively impairs the synthesis of mitochondrial nascent chains. (**A**) A representative 30-minute ^35^S-metabolic pulse labeling of mitochondrial protein synthesis. CBB, coomassie brilliant blue. BMDM, bone marrow derived macrophages. WT, wild type. (**B**) Quantification of nascent chain synthesis for the indicated polypeptides from independent experiments.

**Fig S4:**
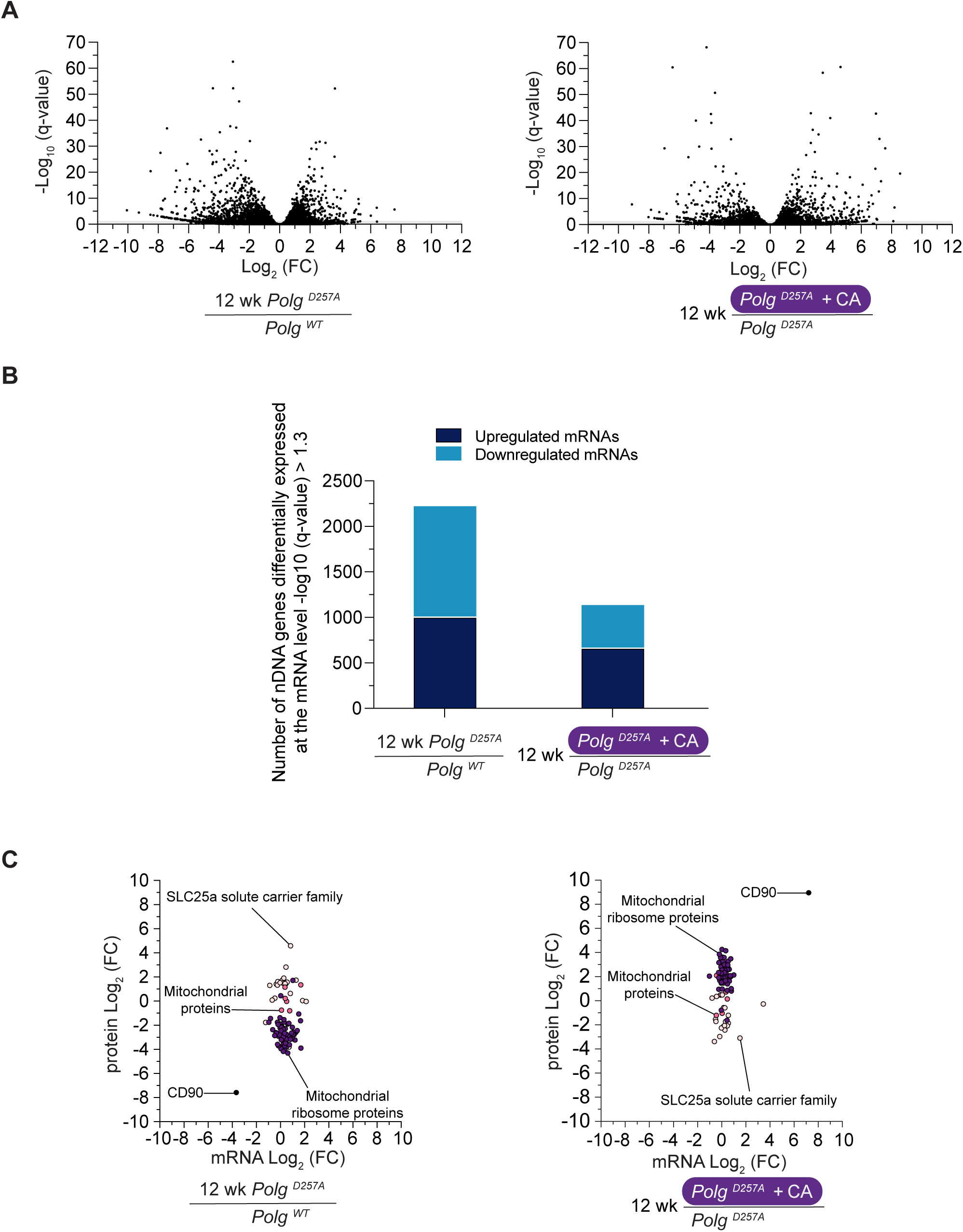
Mitochondrial DNA mutations in *Polg ^D257A^*MEFs significantly alters nuclear gene expression. (**A**) A volcano plot of differential expression from total RNAseq of MEFs comparisons with the indicated genotypes. CA, chloramphenicol. (**B**) Total number of differentially expressed mRNAs q-value < 0.05. (**C**) A scatter plot comparing the log2 expression of mRNA vs. protein abundance of selected factors from the indicated pathways in MEFs.

**Fig S5:**
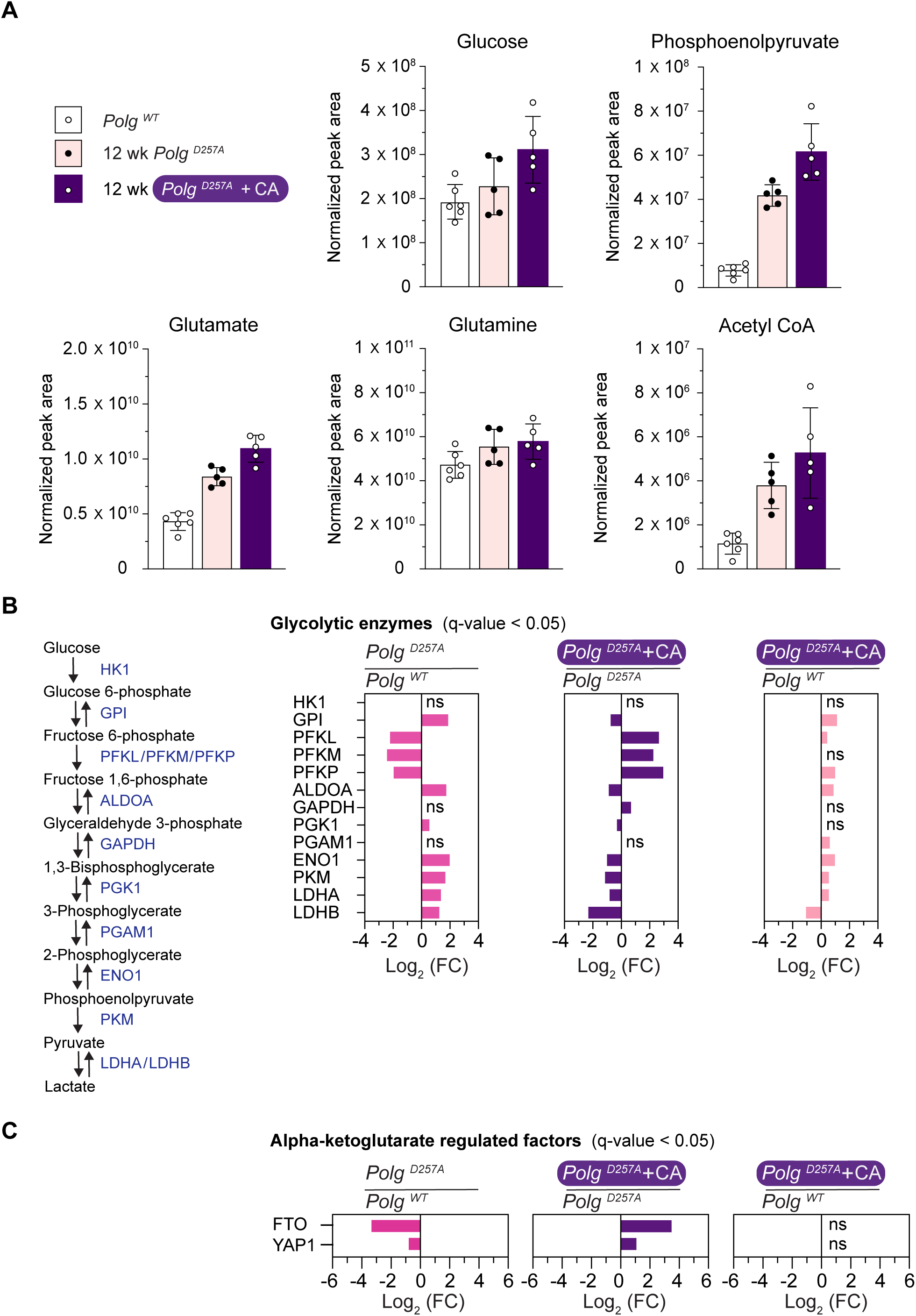
Metabolic rewiring in *Polg ^D257A^* MEFs following inhibition of mitochondrial protein synthesis with chloramphenicol. (**A**) Individual metabolites determined from cells at 12 weeks of culture for *Polg ^D257A^* MEFs treated with and without chloramphenicol compared to wild type *Polg* MEFs. Each data point represents a separate cell sample. Bar graphs are mean +/- SD, n=5. (**B**) Left, a schematic of glycolysis. Right, differential protein abundance of glycolytic enzymes in MEFs with the indicated *Polg* genotypes and treatment conditions determined by label free LC:MS/MS with a q-value < 0.05. ns, non-significant. (**C**) Differential protein abundance of alpha-ketoglutarate dependent RNA demethylase FTO and its substrate YAP1 determined by label free LC:MS/MS with a q-value < 0.05. ns, non-significant.

**Fig. S6:**
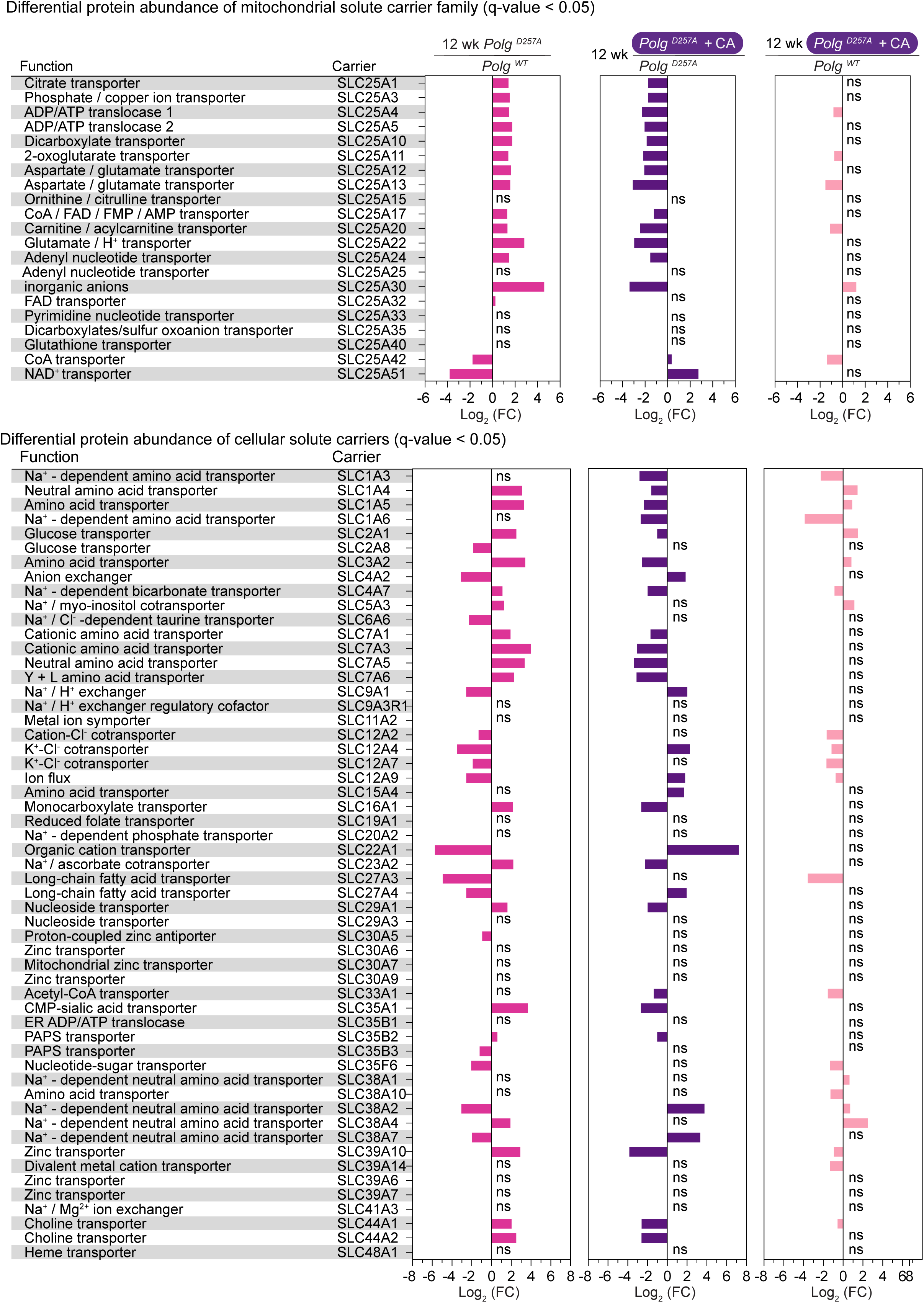
Chloramphenicol inhibition of mitochondrial protein synthesis in *Polg ^D257A^* MEFs significantly modulates the protein abundance of mitochondrial and cellular solute carriers upon long-term culturing. Differential protein abundance in MEFs with the indicated *Polg* genotypes and treatment conditions determined by label free LC:MS/MS with a q-value < 0.05. ns, non-significant.

**Fig. S7:**
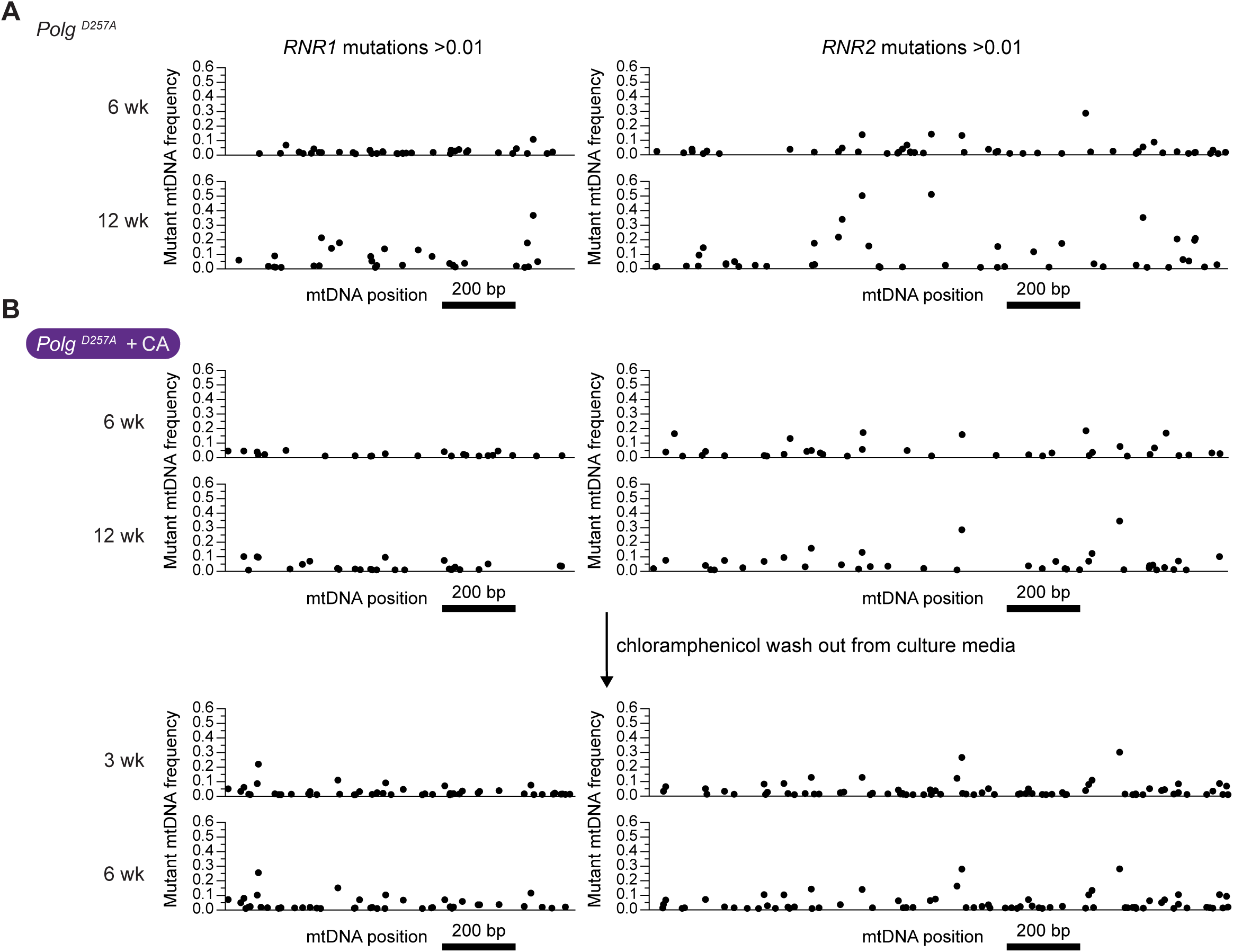
Position of mutations in the mitochondrial rRNA genes from *Polg ^D257A^* MEFs with extended culture. (**A**) *Polg ^D257A^* MEFs following extended culture. (**B**) *Polg ^D257A^* MEFs treated with chloramphenicol during extended culture followed by removal of the antibiotic after 12 weeks for a further 6 weeks of culture.

**Fig. S8:**
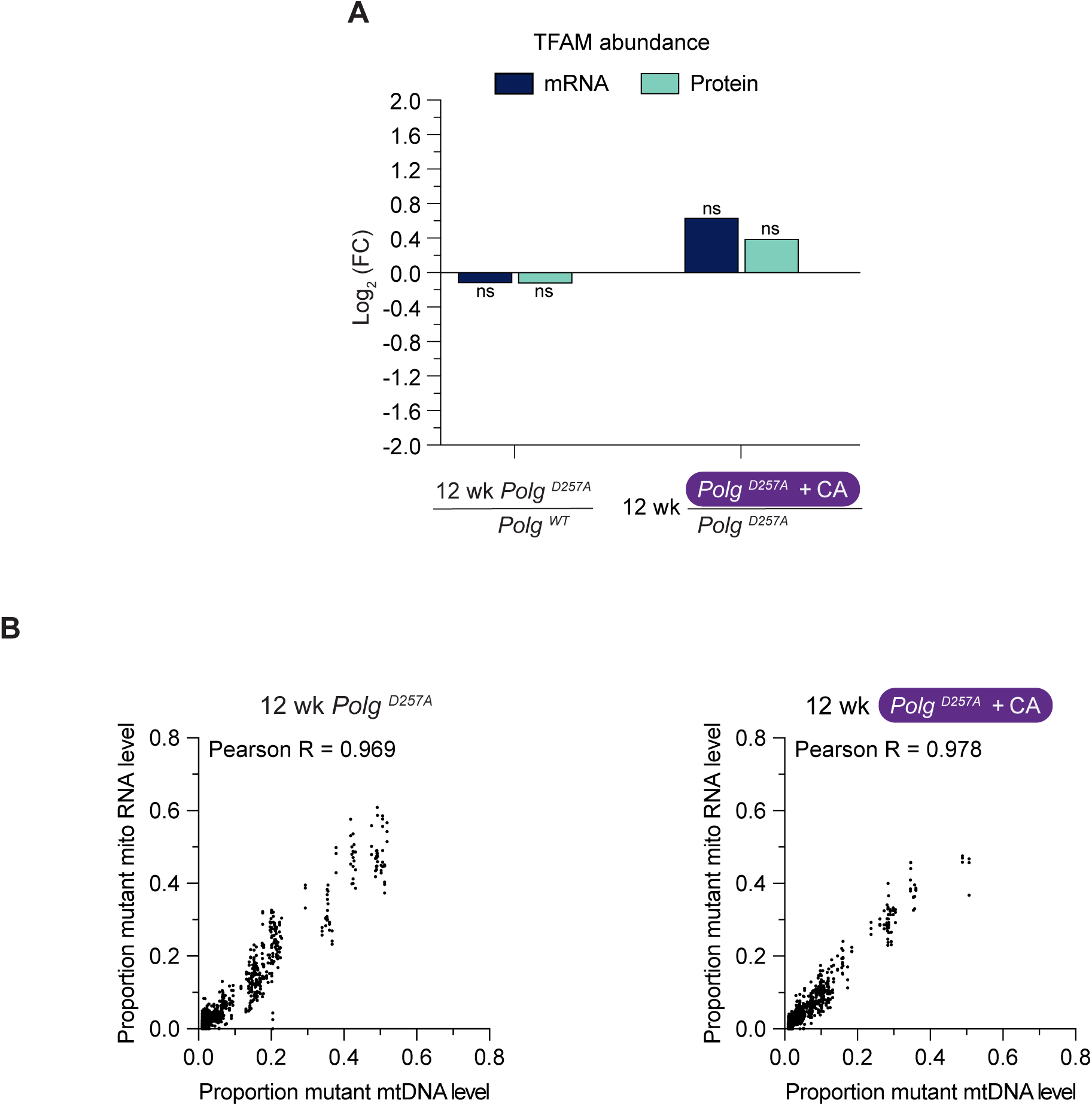
(**A**) **Mitochondrial DNA mutations in *Polg ^D257A^*MEFs expressed at the RNA level.** (**A**) Analysis of TFAM expression at the mRNA and protein level in MEFs. ns, non-significant. (**B**) A scatter plot comparing the heteroplasmy level of a mtDNA mutation expressed at the mitochondrial mtDNA vs. mRNA level (n=3). Pearson correlation coefficient.

**Fig. S9:**
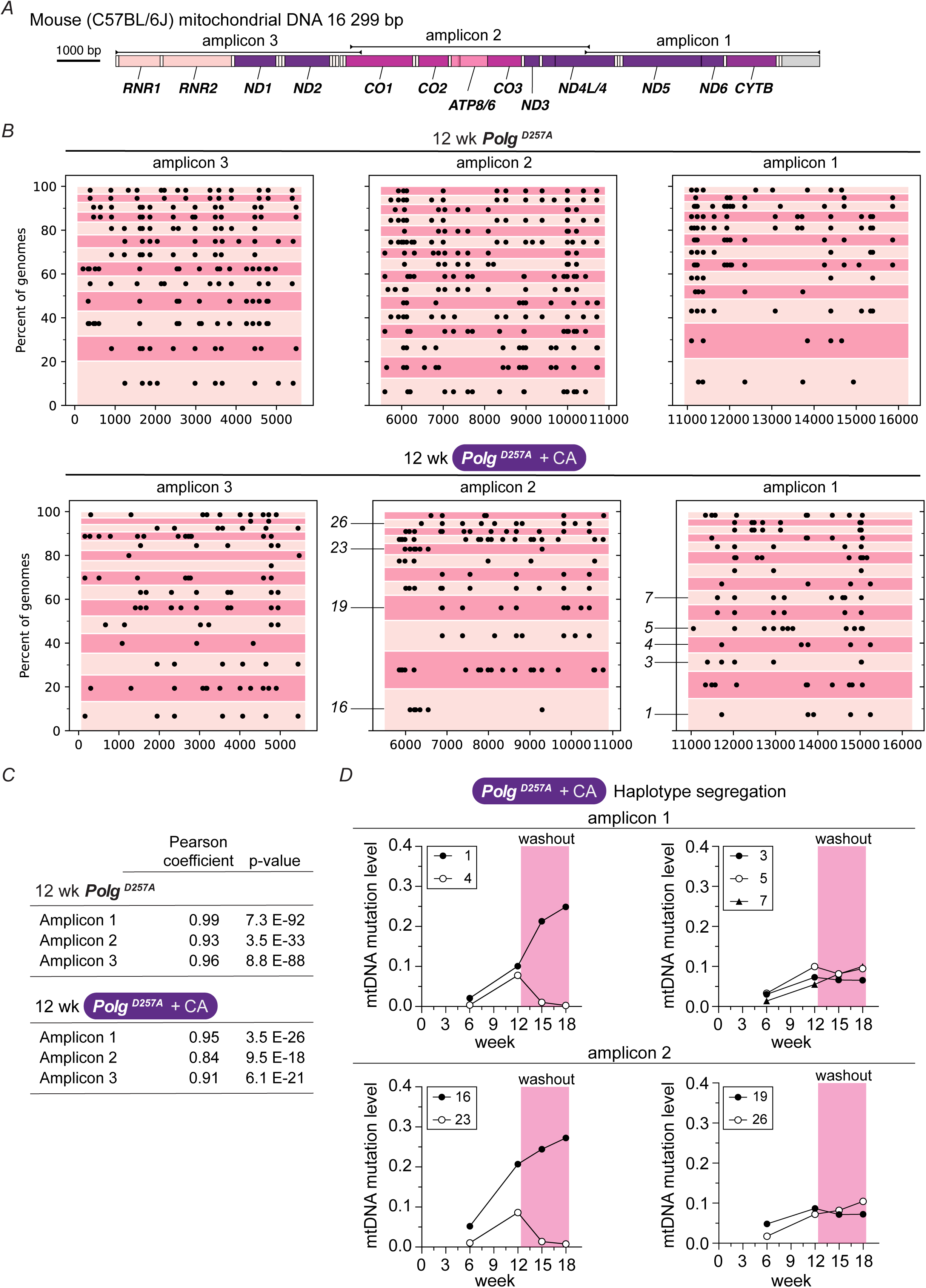
Haplotype segregation of mitochondrial DNA (mtDNA) mutations in *Polg ^D257A^* MEFs with extended culture. (**A**) Schematic of the mouse mtDNA with location of amplicons indicated that were generated for deep sequencing. (**B**) Haplotypes of linked mutations determined by long read PacBio sequencing after 12 weeks of extended culture and their position (black dot) with the x-axis corresponding to the base pair position in the C57BL/6J reference mtDNA genome. The y-axis is a cumulative frequency of the haplotypes. (**C**) Correlation of mtDNA mutation level determined by short read Illumina sequencing of the amplicons vs. long read PacBio. (**D**) Haplotype segregation profile of linked mutations in *Polg ^D257A^* MEFs following extended culture in chloramphenicol and then removal of the antibiotic from the media for a further 6 weeks. The number of an individual haplotype corresponds to those marked in panel (**B**). Mutation level corresponds to the median level of unique mtDNA mutations per haplotype determined by short read Illumina sequencing.

**Fig S10:**
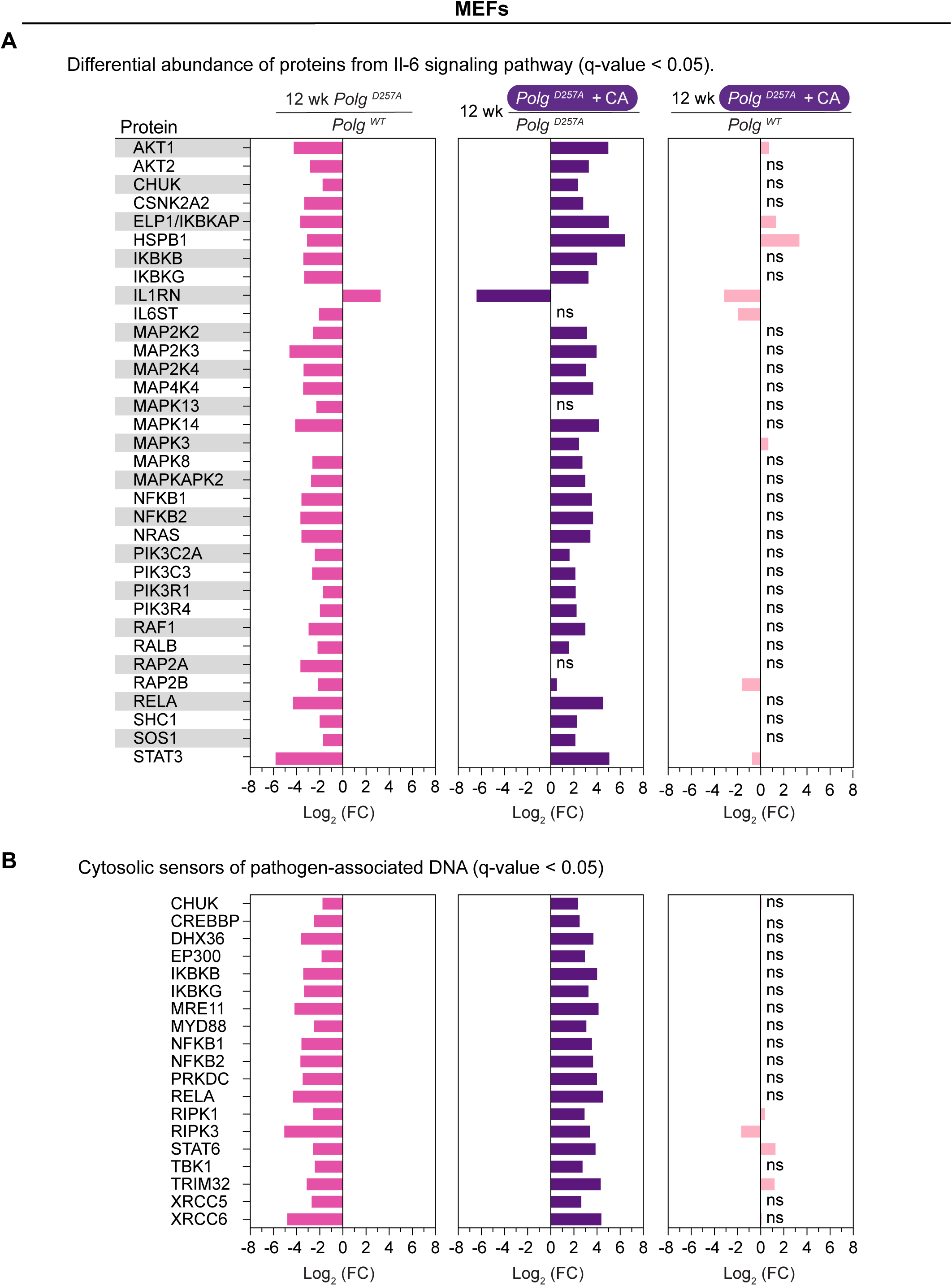
Chloramphenicol inhibition of mitochondrial protein synthesis in *Polg ^D257A^* MEFs reverses the shift in protein abundance of factors in immunity related pathways upon long-term culturing. Differential protein abundance in MEFs with the indicated genotypes and treatment conditions determined by label free LC:MS/MS with a q-value < 0.05. ns, non-significant. The list of proteins determined from the Qiagen Ingenuity Pathway Analysis category of canonical pathways.

**Fig. S11:**
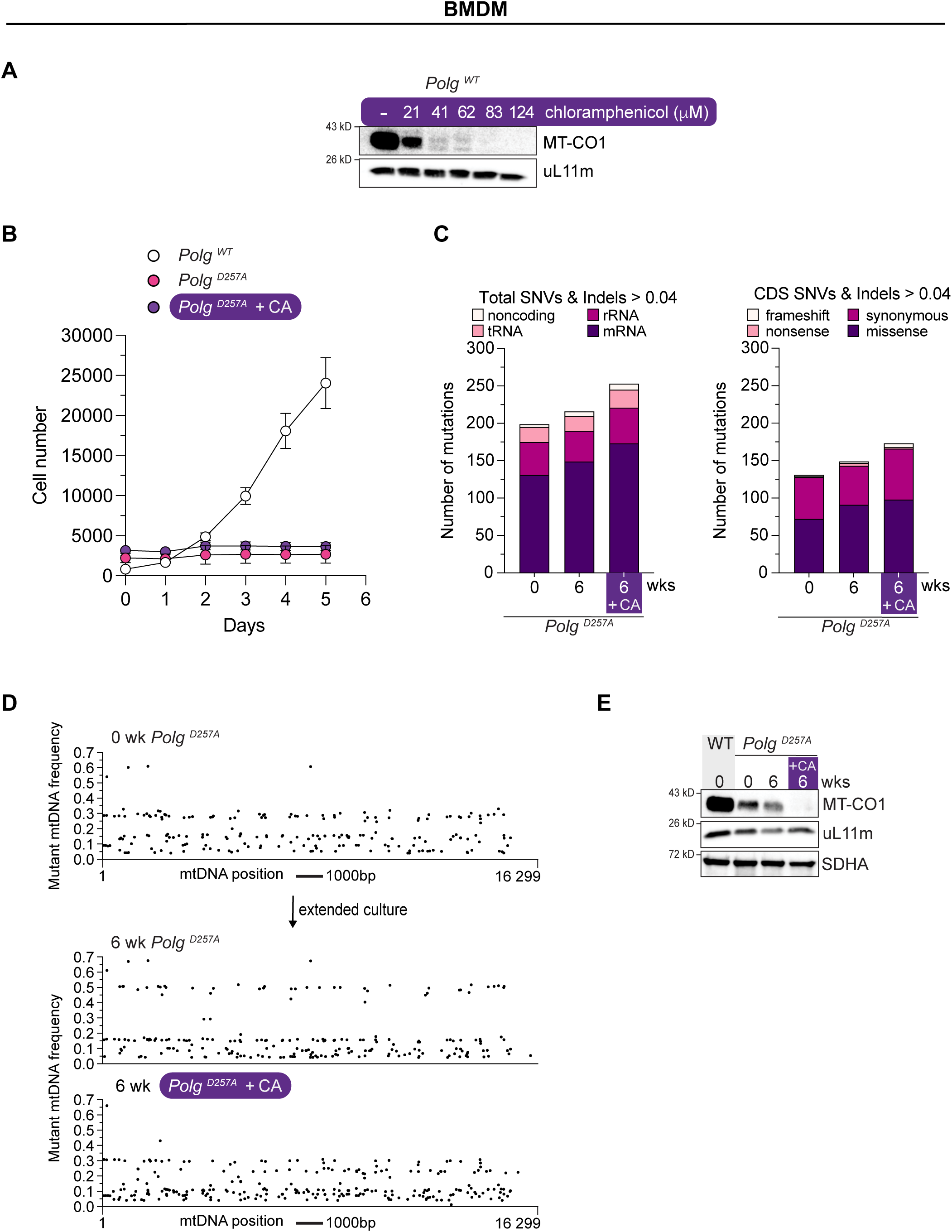
Chloramphenicol inhibition of mitochondrial translation in *Polg ^D257A^* mouse bone marrow derived macrophages (BMDM). (**A**) A chloramphenicol dose-response curve of BMDM with immunoblotting against the indicated proteins after 5 days of treatment. MT-CO1 is encoded in mtDNA. uL11m is a nuclear encoded mitochondrial ribosomal protein of the large subunit. **(B)** Growth curve of early *Polg ^D257A^* BMDM cultures with and without chloramphenicol (CA) over 5 days. Cells were seeded at equal densities and imaged over 5 days by live-cell imaging. (**C**) Mutation analysis of mtDNA isolated from *Polg ^D257A^* BMDMs following extended culture occurring greater than 0.04. (**D**) Distribution of individual mtDNA mutations in *Polg ^D257A^* BMDM along mtDNA following extended culture with and without chloramphenicol in the media. (**E**) Representative immunoblotting of whole cell lysates of bone marrow derived macrophages (BMDM) with the indicated antibodies.

**Fig. S12:**
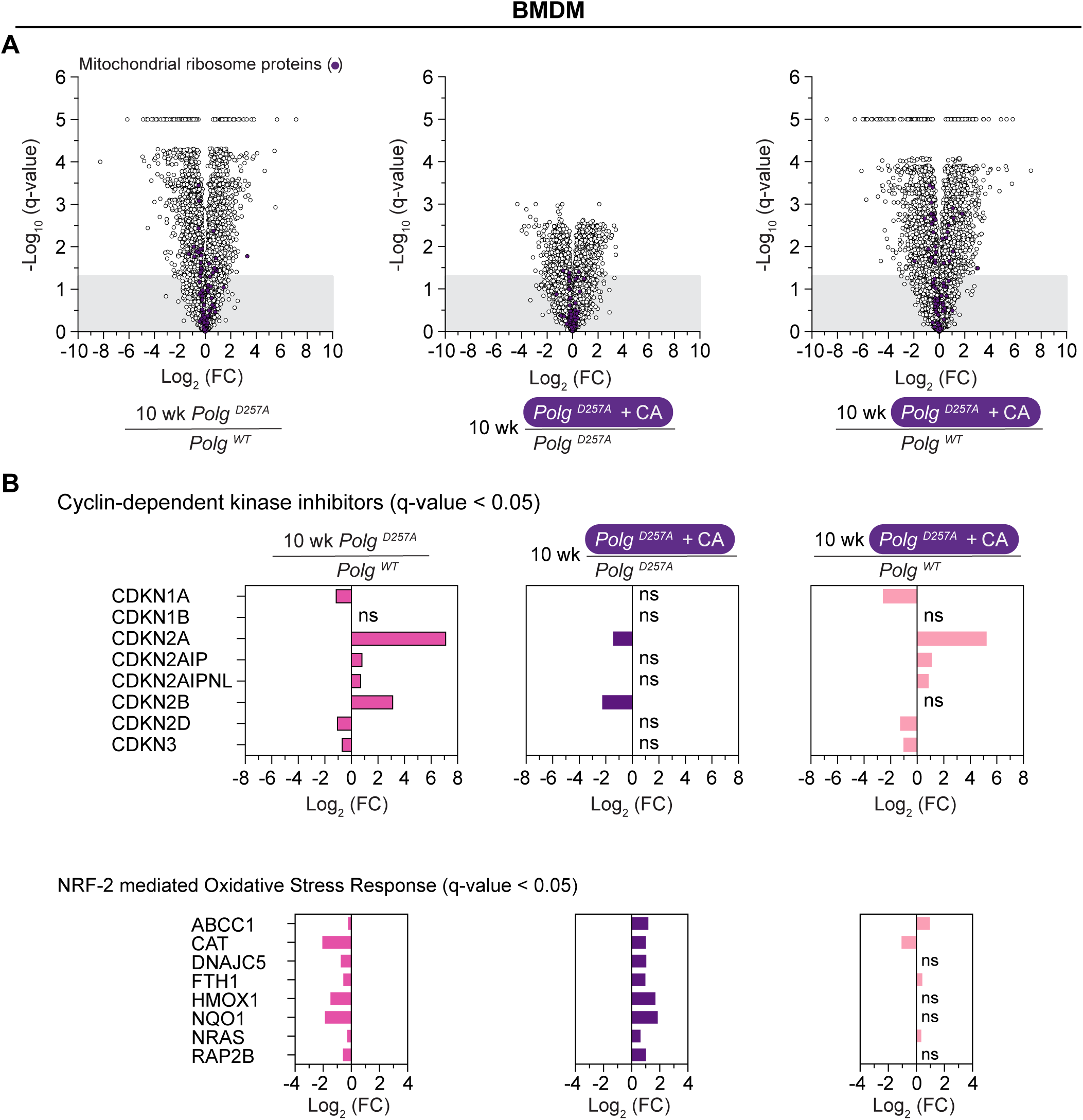
Mitochondrial translation-dependent remodeling of the cellular proteome of *Polg ^D257A^* BMDM following extended culture. (**A**) A volcano plot of protein abundance by quantitative label-free LC-MS/MS for indicated genotypes (n=5) and treatment conditions. (**B**) Differential protein abundance in BMDM with the indicated genotypes and treatment conditions determined by label free LC:MS/MS with a q-value < 0.05. ns, non-significant.

**Fig. S13:**
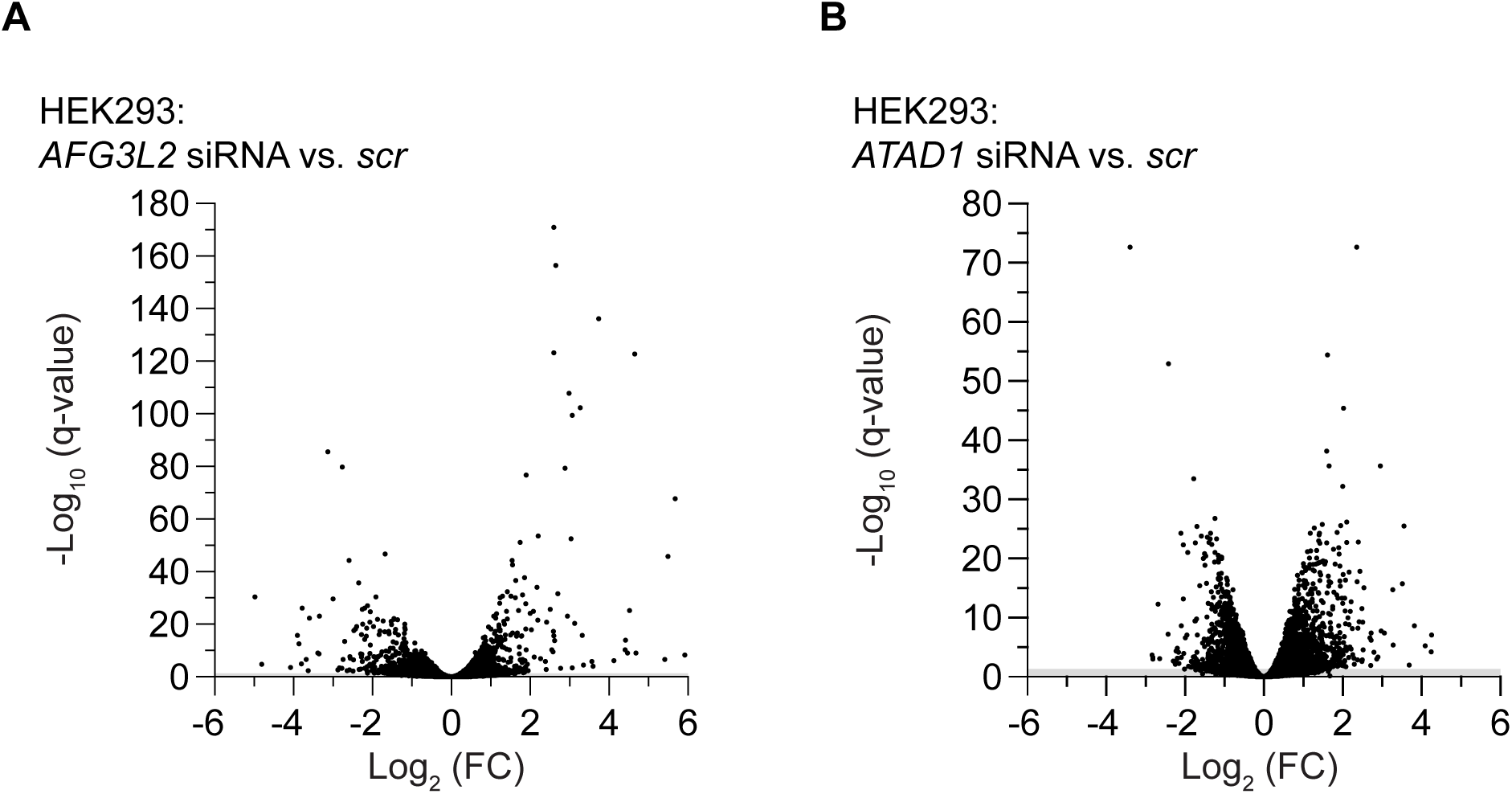
Aberrations in mitochondrial nascent chain synthesis and quality control trigger activation of the integrated stress response and GDF15 expression. Differential expression from total RNAseq. (**A-B**) HEK293 cells treated with small interfering RNA (siRNA) specific to the indicated essential subunits of mitochondrial AAA protein complexes anchored in the mitochondrial inner membrane (AFG3L2) or outer membrane (ATAD1).

